# Brain network reconfiguration during prediction error processing

**DOI:** 10.1101/2023.07.14.549018

**Authors:** Kamil Bonna, Oliver James Hulme, David Meder, Włodzisław Duch, Karolina Finc

## Abstract

Learning from experience is driven by reward prediction errors—signals that reflect updates to our expectations of reward. Despite numerous studies on neural correlates of reward prediction errors, the question of how large-scale brain networks reconfigure in response to reward prediction error signalling remains open. Here we ask how functional networks change in response to reward prediction errors depending on the context. In our study participants performed the probabilistic reversal learning task in functional magnetic resonance imaging (fMRI) scanner in two experimental contexts: a reward-seeking setting and a punishment-avoiding. We found that the participants’ learning speed depended on the sign of the prediction error but not on the experimental context. Whole-brain network analysis revealed a multi-scale community structure with a separate striatal reward network emerging at a finer topological scale and a ventromedial prefrontal network emerging at a coarser scale. We also found that the integration between large-scale networks increased when switching from positive to negative prediction error events. This pattern of large-scale network reconfiguration aligns with the broad range of research showing increased network integration with increased cognitive demands. Our findings offer a first sketch of how processing reward prediction error affects the functional connectivity of brain-wide networks.

## Introduction

Learning through interaction with the environment is an essential feature of all intelligent systems. In the context of *reinforcement learning*, where rewarding events reinforce behavior and punishing events have the opposite effect (Sutton & Barto, 2018), prediction error coding plays a critical role. This mechanism continuously updates future expectations of rewards based on the disparities between predicted and actual outcomes (Rao & Ballard, 1999). Remarkably, neuroscience research has demonstrated that the brain encodes reward prediction errors by modulating the phasic activity of dopaminergic neurons (Colombo, 2014; Montague *et al*., 1996), which project throughout the entire brain, engaging interconnected networks (Jaber *et al*., 1999). Despite extensive investigations into the neural correlates of reward prediction errors, the impact of such coding on the entire brain network remains an intriguing and unanswered question.

One of the crucial attributes of the reward prediction error is its *sign*. The sign of the reward prediction error, distinguishing between better-than-expected and worse-than-expected outcomes, is used to differentially value particular actions, policies, and beliefs that contributed to the outcome. The better-than-expected outcomes cause an increase in the assigned value, whereas worse-than-expected outcomes cause a decrease of current value. From a computational point of view, positive and negative errors arise from the same underlying computation that estimates reward value functions, mapping rewarding and punishing events onto a common scale, ranging from negative to positive. From this perspective, a single neural circuit could be sufficient for computing both positive and negative prediction errors. Yet, multiple studies report that brain areas involved in negative prediction error processing are located also outside the dopaminergic system (Fazeli & Büchel, 2018; Hauser *et al*., 2015; Yacubian *et al*., 2006).

Palminteri and Pessiglione (2017) suggested the existence of an opponent system carrying a negative portion of the prediction error signal. Consistent with this idea, a recent meta-analysis identified two spatially distinct learning networks for processing positive and negative prediction errors (Fouragnan *et al*., 2018). The network coding positive prediction errors involved regions such as ventrolateral prefrontal cortex (vlPFC), ventral striatum (VS), and posterior cingulate cortex (PCC), while the network coding negative prediction errors involved regions such as dorsomedial cingulate cortex (dMCC), anterior insula (aINS), pallidum, and middle frontal gyrus (MFG). Despite many studies focusing on the neural underpinnings of positive and negative prediction errors, research into the whole-brain interactions associated with the prediction error processing is lacking. It is still unclear how the brain systems coding positive and negative prediction errors might interact between themselves and with other subsystems of the brain.

There is some evidence that the sign of the reward prediction error depends not only on the decision outcome, but is also modulated by the environmental context in which these errors occur (Bavard *et al*., 2018; Rangel & Clithero, 2012). Positive prediction errors are usually related to rewarding outcomes, but they can also signal relief from avoiding punishment when the agent perceives the environment as generally adverse (Nieuwenhuis *et al*., 2005; Palminteri *et al*., 2015). Equivalently, negative prediction errors can arise when the reward is anticipated but not provided in a rewarding environment. Such context-dependence may suggest that the brain uses a *reference point* to which experienced outcomes are compared (Bavard *et al.*, 2018; Rangel & Clithero, 2012). The reference point hypothesis states that decision values are actively constructed based on average values present in the environment. It enables utilizing both positive and negative prediction error systems regardless of the distribution of rewards and punishments. The reference hypothesis was supported by both behavioral (Khaw *et al*., 2017; Louie *et al*., 2013) and neuroimaging studies using fMRI (Park *et al*., 2012; Rigoli, 2019). Using outcome valence as an explicit experimental factor, studies have found that the ventral reward system signaled positive prediction errors irrespective of the outcome valence (Meder *et al*., 2016; Palminteri *et al*., 2015). On the other hand, the amygdala, inferior frontal gyrus (IFG), and dorsomedial prefrontal cortex (dmPFC) signaled positive prediction errors in reward but not in the punishment context (Meder *et al*., 2016). Also, Robinson *et al*. (2010) demonstrated simultaneous valence-specific and valence-nonspecific signals in the stiatum. Due to these inconclusive findings, context-dependence of prediction error coding is still a debated issue. Moreover, it is unknown whether the reference effect is also reflected by the outcome valence invariance at the whole-brain network level.

In recent years, the interdisciplinary field of *network neuroscience* (Bassett & Sporns, 2017) has provided valuable insights into the dynamic nature of functional brain networks, highlighting their capacity for reorganization in response to environmental demands (Bullmore & Sporns, 2009). Notably, fluctuations of the modular structure of the brain network across temporal scales, have been observed across various cognitive processes and tasks (Vatansever *et al*., 2015, Finc *et al*., 2017, Betzel *et al*., 2018; Shine *et al*., 2019). To date, there have been few attempts to investigate large-scale brain networks during reinforcement learning. Most studies investigated gradual network changes associated with reward learning, not focusing on rapid changes related to switching between positive and negative prediction errors (Gerraty *et al*., 2018; Mattar *et al*., 2018). Sadler *et al*. (2020) used a taste-motivated learning task to examine the dynamics of the reward network and found that the community structure of the functional brain networks was affected by the prediction error sign—ventromedial and ventrolateral prefrontal cortices changed their module assignment during sweet taste eliciting positive prediction errors compared with bitter taste eliciting negative prediction errors. This limited evidence leaves the relationship between whole-brain network dynamics and prediction error coding largely unexplored.

A few studies have investigated context-dependence (reward vs. punishment) of connectivity profiles during prediction error processing. In one study, Camara *et al*. (2009) investigated functional connectivity patterns during monetary gain and losses. For example, network of brain regions encompassing the amygdala, orbitofrontal cortex, and insula was coupled with the VS during monetary gambling irrespectively of the outcome valence. However, the orbitofrontal cortex had stronger connections with VS during punishing trials. Other studies found increased functional connectivity between VS and ventromedial prefrontal cortex and between the amygdala and VS, midbrain, cingulate cortex, thalamus, orbitofrontal cortex, and dorsolateral prefrontal cortex during reward processing (Van den Bos *et al*., 2012, Cohen *et al*., 2005). Overall, these findings yield inconsistent answers to the question of whether regions encoding prediction errors share similar or different connectivity profiles during reward and punishment processing.

The main goal of this work is to provide a comprehensive description of the large-scale network reconfiguration during prediction error processing in reward and punishment contexts. We asked 32 healthy participants to play a probabilistic reversal learning paradigm in both reward-seeking and punishment-avoiding conditions, whilst acquiring whole brain fMRI data. First, we modeled participants’ behavior with a Bayesian model with four competing submodels and assessed which computational model of reinforcement learning best explains the subjects’ decisions. Then, we used parameters of the best fitting model to investigate how the organization of the brain network shifts when participants switch between the processing of positive and negative prediction errors. As the functional brain network is known to exhibit a multiscale community structure (Betzel & Bassett, 2017), we have focused on describing modular network reorganization on multiple topological scales. This allowed us for for capturing effects that might have been overlooked in a single-scale analysis (Betzel *et al*., 2015). Here we hypothesized that (1) switching between positive and negative prediction errors will lead to dynamic reorganization of the brain network and the formation of prediction-error-specific subnetworks, (2) regions signaling prediction errors will form separate subnetworks that should be detectable for finer topological scales, and (3) prediction-error-specific subnetworks will interact with other large-scale networks involved in cognitive task processing, especially the default mode network (DMN), and the fronto-parietal network.

In the reversal learning task, positive prediction errors usually confirm the subject’s internal judgment about a more beneficial option and are typically followed by choice repetition. On the other hand, negative prediction errors typically lead to a conflict—the subject has to judge whether the source of the error lies in the environmental stochasticity or reflects a genuine change in the causal structure of the environment. This conflict plausibly requires the increased engagement of cognitive resources as a new environmental structure is entertained. For example, Finc *et al*. (2017) found cognitive load-related modularity breakdown resulted from decreased segregation of the DMN and its increased integration with other large-scale networks, especially task-positive ones. Here, we hypothesized that we will observe the the same effect of network switching from positive to negative reward prediction errors. Specifically, we expected to observe decrease of segregation of the DMN and the reward network, and increase of integration between these networks and the task-positive networks. We also hypothesized that the whole-brain network modularity will decrease along with decreasing reward prediction errors. Lastly, we expected to find support for a reference point hypothesis, which states that prediction error systems should be invariant to the outcome valence. On the behavioral level, this hypothesis predicts that the subject’s learning rates for positive and negative prediction errors are invariant to the task condition. On the level of brain networks, it might suggest that neural correlates of prediction errors should be identical during reward-seeking and punishment-avoiding conditions.

## Results

### Behavioral modeling of prediction error coding in reward-seeking and punishment-avoiding conditions

To study prediction error coding in different contexts, we used a *probabilistic reversal learning* (PRL) task (**Fig. 1**; Cools *et al*., 2002). The PRL task is a powerful paradigm for investigating the process of updating and relearning the stimulus-reward association through reinforcement. Here, participants performed the PRL task in two conditions: reward-seeking (RS) and punishment-avoiding (PA). In each condition winning and losing are opposed to neutral outcomes, enabling disentangling between often confused dimensions: outcome valence and prediction error sign (Palminteri & Pessiglione, 2017). This quality is crucial to investigate how these dimensions independently influence decision behavior and neural correlates of prediction errors.

**Fig. 1.**
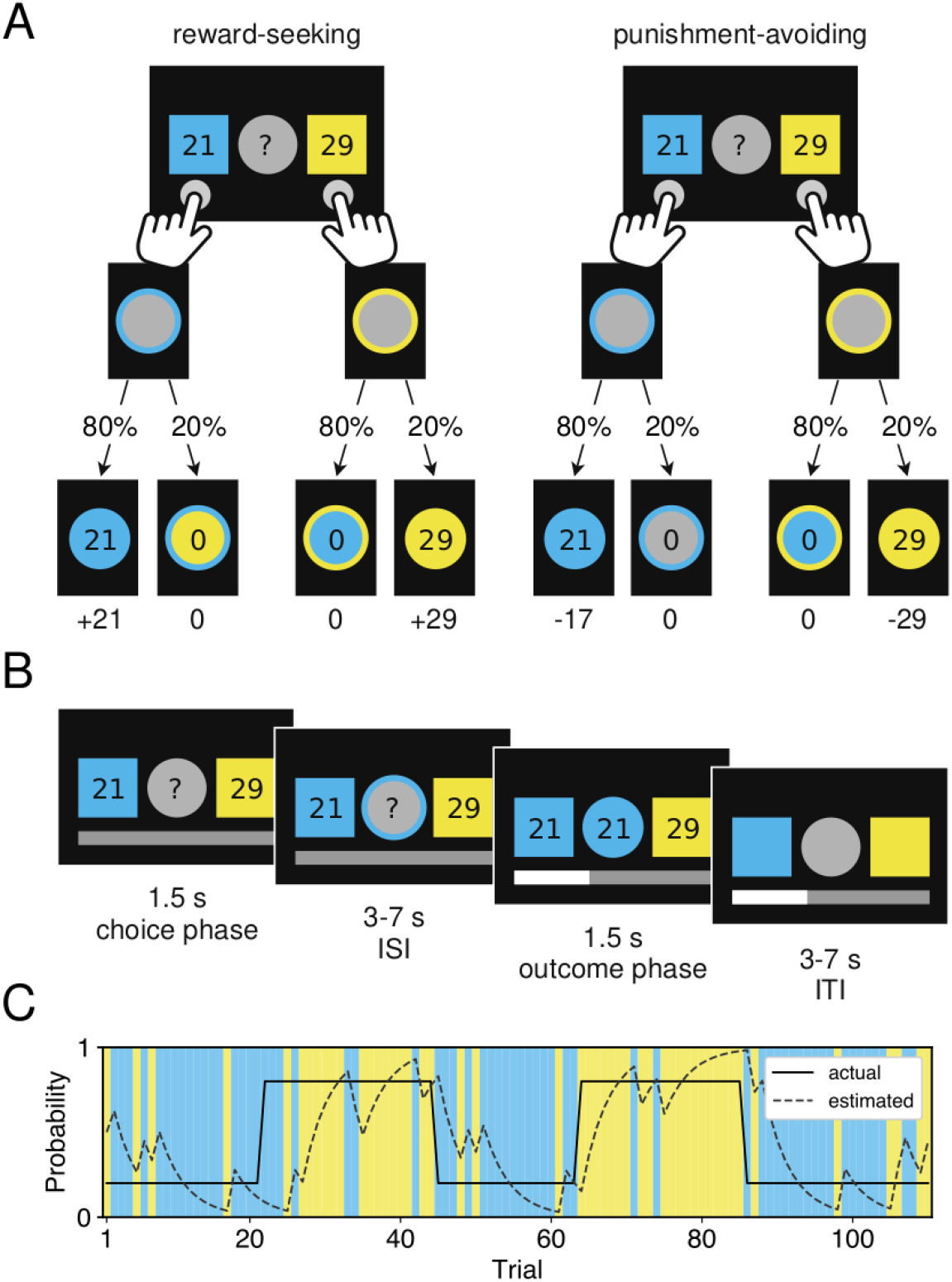
Probabilistic reversal learning task. (**A**) Task structure for reward-seeking and punishment-avoiding conditions of the PRL task, represented as a decision tree. In the punishment-avoiding condition, numbers in the boxes represented punishment magnitudes, the fixation circle changed its color according to the punished box, and the number of lost points was displayed in the middle. (**B**) Example trial of probabilistic reversal learning task in reward-seeking condition. Each trial began with a choice phase where the subject had 1.5s to choose between blue and yellow boxes. Subjects had to consider both reward magnitudes visible on the boxes and reward probabilities estimated from previous trials. After variable inter-stimulus-interval (ISI), an outcome phase began, giving the subject choice feedback. During the outcome phase, the fixation circle changed its color according to the rewarded box, and the number of rewarded points was displayed in the middle. The outcome phase was followed by a variable inter-trial-interval, after which a new trial began. (**C**) Each task condition consisted of 110 trials. Reward contingency changed four times throughout the task. The solid line represents true reward probability for the yellow box, whereas the dashed line represents prediction of a standard TD model.

An individual subject’s performance accuracy was calculated to establish if subjects succeeded in learning probabilistic associations. Accuracy was defined as the proportion of choices that led to rewarding outcomes in the reward-seeking condition or non-punishing outcomes in the punishment-avoiding condition. Since the study examined healthy subjects without valence-dependent learning deficits, we hypothesized similar performance levels for both task conditions.

The accuracy was significantly higher than chance level for both reward-seeking (acc_RS_ = 62.81%; one-sided t-test *t*(31) = 13.00; *p* < 0.0001) and punishment-avoiding condition (acc_PA_ = 62.13%; one-sided t-test *t*(31) = 11.38; *p* < 0.0001). Subjects performed equally well in both experimental contexts—no significant difference in performance between task conditions was found (two-sample t-test; *p* = 0.62). Moreover, the number of choice reversals was calculated for each subject and task condition. Subjects frequently reversed their choices—mean number of reversals was equal to 30.41 ± 11.93 in the reward-seeking and 30.25 ± 11.98 in the punishment-avoiding condition. Moreover, the number of choice reversals did not significantly differ between the two task conditions (two-sample t-test; *p* = 0.94).

Finally, we quantified differences in choice duration, i.e., reaction time (RT), between reward-seeking and punishment-avoiding conditions, and between positive and negative prediction errors. Choice times following gain/no-lose trials were assigned to the +PE group, whereas these following lose/no-gain trials to the −PEs. A two-way repeated measures ANOVA (rmANOVA) was performed on RT values with task condition and PE sign as factors. There was a significant interaction effect between task condition and PE sign (*F*(31) = 14.06; p < 0.001; **Fig. A.1B**) indicating that choice durations were modulated by both experimental factors. In general, choice durations were longer for the punishment-avoiding compared to reward-seeking conditions, as indicated by the significant task condition effect (RT_RS_ = 687.4 ± 126.3ms; RT_PA_ = 651.7 ± 110.9ms; *F*(31) = 4.27; p < 0.05). Post hoc within-condition analysis showed that choice times following negative prediction errors were longer in punishment-avoiding condition (paired t-test; *t*(31) = 3.04; *p* < 0.01) but not in reward-seeking condition (paired t-test; *p* = 0.26).

Next, a Bayesian hierarchical latent-mixture (HLM) model was used to perform model selection and parameter estimation (see **Supplementary Information 1.1**). This type of Bayesian model contains parameters representing behavioral parameters, like learning rates or precision, and parameters reflecting the belief about which competing model generates the observed data. Following previous studies, models with separate learning rates for positive and negative prediction errors were considered to account for possible risk-sensitivity effects. These models would be referred to as prediction-error dependent (PD) models (Niv *et al*., 2012; Reiter *et al*., 2016; Van den Bos *et al*., 2012). To capture possible valence effects, models with separate learning rates for reward-seeking and punishment-avoiding conditions were also included. These models would be referred to as condition-dependent models (CD). To fully explore model space, all possible combinations of models were considered, which resulted in four different models: (1) PICI model with single learning rate independent of the sign of prediction error and outcome valence, (2) PICD model with two separate learning rates for reward-seeking and punishment-avoiding conditions, (3) PDCI model with two separate learning rates for positive and negative PEs, (4) PDCD model with four separate learning rates for prediction error signs and task conditions.

The HLM model revealed that a majority of the subjects (27/32; 84.4%) had most of the probability mass located over either the PDCI model (19/32; 59.4%) or PDCD model (8/32; 25%) (**Fig. S2A**). Models with separate learning rates for reward-seeking and punishment-avoiding conditions were favored in five subjects (1/32; 3.1%; PICD and 4/32; 12.5% PICI). Posterior distribution of the model indicator variable was marginalized over subjects and model classes to investigate whether subject response patterns are better explained by models with: (1) prediction error dependent (PD) versus prediction error independent (PI) learning rates, and (2) condition dependent (CD) versus condition independent (CI) learning rates. These comparisons reflected two orthogonal experimental axes: prediction error sign and task condition. Moderate evidence was found in favor of the hypothesis that behavioral responses are better explained by the models with separate learning rates for positive and negative prediction errors (BF_PD−PI_ = 3.19; **Fig. S2B**). On the other hand, there was weak evidence against the hypothesis assuming separate learning rates for reward-seeking and punishment-avoiding conditions (BF_CD−CI_ = 0.67; **Fig. S2C**). Measures of the likelihood for each submodel being more frequent than all other submodels of the HLM model were computed as protected exceedance probabilities. This procedure determined a single winning submodel for further fMRI analysis. The PDCI model was the most likely model across participants with probability close to 1 (protected exceedance probability; *p* = 0.9995; **Fig. S2D**).

### Functional networks involved in prediction error processing

Recent research identified two brain systems responsible for coding positive and negative prediction errors (Fouragnan *et al*. 2018). How do these two brain systems interact between themselves and with other subsystems during reinforcement learning in reward-seeking and punishment-avoiding conditions?

First, we applied standard general linear model (GLM) analysis (see **Supplementary Information 2.2**), to investigate activation pattern during processing positive and negative prediction errors during differend contexts. We identified two sets of regions associated with positive and negative prediction error processing, which were consistent with previous findings from the literature (Meder *et al*., 2016, see **Fig. S4** and **Tab. S1**). Bilateral visual areas V3 and V4, right supramarginal gyrus, right superior parietal lobule, right precuneus, right primary visual cortex, and right precentral gyrus, showed higher activity during reward-seeking compared to punishment-avoiding condition.

Then, to investigate how brain regions coding positive and negative prediction errors interact with other subsystems at the network level, we used beta-series correlation analysis. First, we extended a standard parcellation introduced by Power *et al*. (2011), including 264 regions of interests (ROIs), to include brain regions identified in a recent meta-analysis on neural representations of prediction error valence (Fouragnan *et al*., 2018) (**Fig. 2**, **Supplementary information 2.3.1**). This procedure allowed us to study effects specific for networks signaling prediction errors while still preserving reference to the well-known large-scale brain systems. Afterwards, we used parameters of the best fitting model to investigate how the organization of the brain network shifts when participants switch between the processing of positive and negative prediction errors, testing the hypothesis that two separate networks are responsible for positive and negative prediction error processing.

**Fig. 2.**
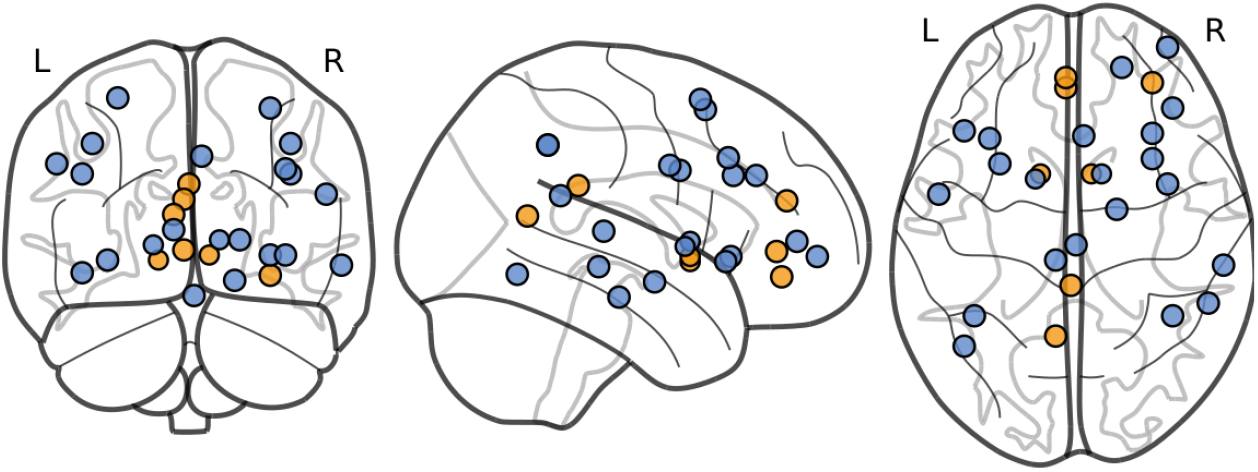
Prediction error signaling ROIs. Regions of interest created from activation likelihood estimation (ALE) meta-analysis clusters from Fouragnan *et al*. (2018). Network signaling positive prediction errors (+PE; orange spheres) is created from regions with higher BOLD activation for positive than negative PEs. Network signaling negative prediction errors (−PE; blue spheres) is created from regions with higher BOLD activation for negative than positive PEs.

Iinstead of only testing for condition-related regional activity changes, exploring the pattern of changes in modular network organization can provide additional insights into how the brain achieves learning and decision-making in different contexts. Here we used an approach based on within- and between-community agreement to investigate whether the integration and segregation of large-scale networks depend on the prediction error sign and outcome valence. This community-level approach relies on averaging node-level agreement, *D_ij_*, derived from data-driven partitions using a reference partition. The reference partition can be either a consensus partition, or an apriori partition representing a stable community structure (Conrad *et al*., 2020). To better reflect the underlying structure of the data, the condition-independent consensus partitions were used as reference partitions (see **Fig. 3**). Community-level agreement values capture the level of association between large-scale networks (LSNs); when calculated for a single partition, they represent a subject-level pattern of integration and segregation of different brain systems.

**Fig. 3.**
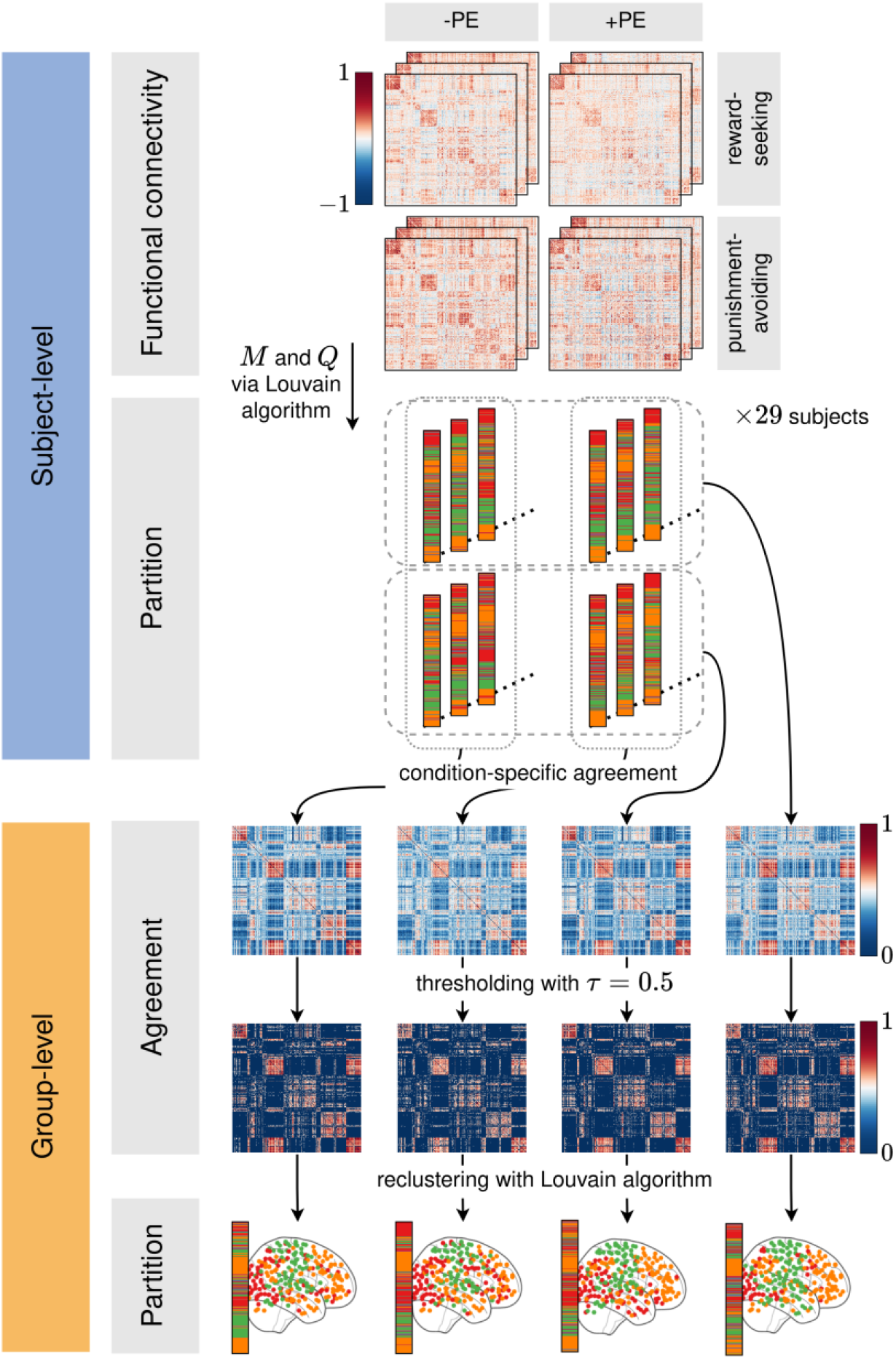
Consensus partitioning pipeline. Multi-step procedure employed to calculate representative network partitions for task conditions and prediction error signs. Subject-level connectivity matrices were first clustered via the Louvain algorithm to calculate network modularity, *Q*, and community affiliation vectors, *M* (Blondel *et al*., 2008). Then, subject-level partitions were grouped into four groups: reward-seeking, punishment-avoiding, positive PE, and negative PE. For each group, the agreement matrix was calculated, thresholded and reclustered, yielding a final condition-specific consensus partition.

*Within-community agreement* expresses the probability that two randomly selected nodes from a given LSN will be found within the same data-driven community. For example, a within-community agreement of 1 would reflect that all LSNs nodes are a part of the same data-driven community. High values of within-community agreement indicate stable and segregated LSN, whereas lower values characterize more unstable and fragmented systems. *Between-community agreement* reflects the probability that a randomly selected pair of nodes from two different LSNs will share the same data-driven community. In most extreme cases, the between-community agreement can be 0 when all nodes from both LSNs belong to different data-driven communities, or 1 when both LSNs are merged into a larger community in a data-driven partition. Between-community agreement expresses the extent of integration between LSNs.

Then, we investigated the modularity of the functional network across three topological scales controlled by *γ* resolution parameters (see **Methods** and **Supplementary Information** for details). On the level of whole-brain network topology, we were interested in whether the overall degree of modularity changes when subjects switch between processing (1) positive and negative prediction errors in (2) risk-seeking and punishment-avoiding environments. We performed two-way repeated measures ANOVA (rmANOVA) on modularity values with two factors: prediction error sign and task condition. The same statistical testing was applied to all three topological scales. Average network modularity was highest for the largest topological scale (*Q*(*γ* = 0.5) = 0.48 ± 0.03) and smallest for the finest topological scale (*Q*(*γ* = 0.5) = 0.16 ± 0.07). It decreased significantly with increasing structural resolution parameter (*p* < 10 −10; paired t-tests between scales). However, the rmANOVA results were not significant. All three effects: main prediction error sign effect, main task condition effect, and interaction (prediction error sign × task condition) effect, did not reach the significance threshold regardless of the topological scale (**Fig. 5**). This shows that the degree of whole-brain network segregation expressed as modularity is stable across task conditions and topological scales.

**Fig. 4.**
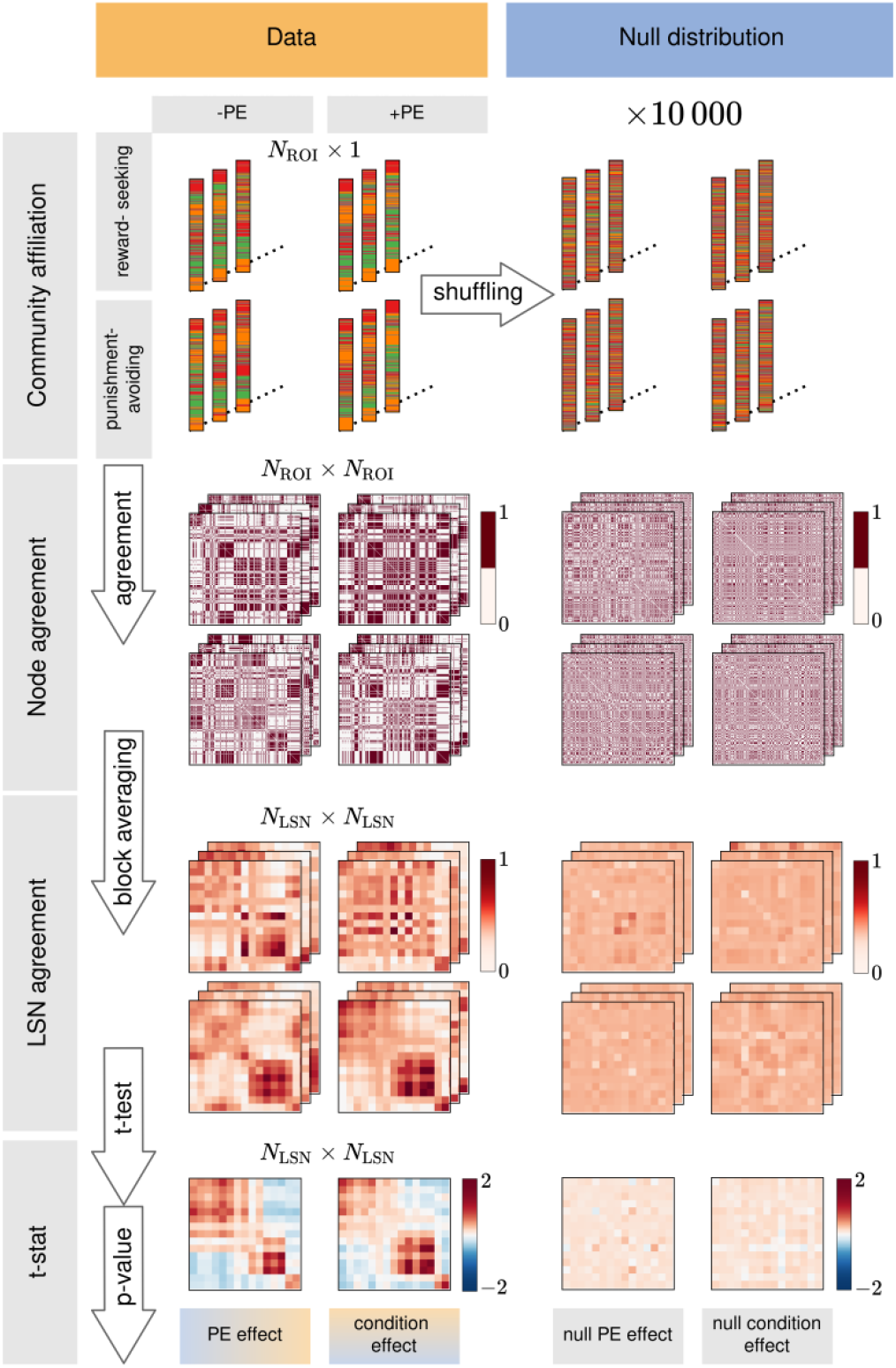
Large-scale network agreement pipeline. Multi-step procedure was employed to calculate condition-specific changes in the community-level agreement. Subject-level community affiliation vectors were used to calculate node-level agreement matrices, which were block averaged to produce individual LSN agreement matrices. LSN agreement represents the probability that two randomly selected nodes from reference LSNs belong to the same data-driven community. For each entry in the LSN agreement matrix, two t-tests were conducted, testing the effect of task-condition and prediction error sign. T-test significance was tested using the Monte Carlo permutation procedure. Shuffled community affiliation vectors were randomly reordered 10,000 times and subjected to the same analysis procedure as the original vectors to produce a null distribution for both t-statistics. P-values were assigned according to the position of the true t-statistic within the null distribution.

**Fig. 5.**
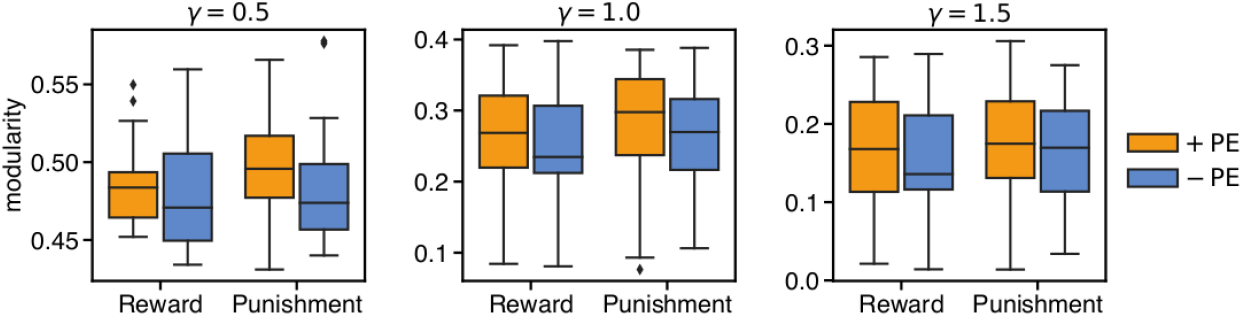
Whole-brain network modularity. Network modularity values for different topological scales and task-conditions. Each plot corresponds to a different topological scale characterized by structural resolution parameter *γ*. Modularity values are grouped according to prediction error sign (orange bars - positive prediction errors; blue bars - negative prediction errors) and task conditions (Reward - reward-seeking condition; Punishment - punishment-avoiding condition). The boxes show the quartiles of modularity distribution while the error bars show lower and upper limits.

Since the same degree of segregation can characterize a large diversity of network architectures, it is important to analyze networks on finer organizational scales of communities and individual nodes. Here, we were interested in whether large-scale network organization differs between reward-seeking and punishment-avoiding conditions and positive and negative prediction errors. We calculated consensus network partitions for each experimental condition and topological scale. We also calculated condition-independent consensus partitions representing stable community structure during prediction error processing. Condition-independent partitions were further used as reference partitions to test condition-specific changes in large-scale networks agreement.

As expected, the number of communities increased, and the average community size for condition-specific consensus partitions decreased with increasing structural resolution parameter *γ*. For *γ* = 0.5, consensus partitions consisted of few “super-communities” (average 2.25 communities across conditions). The mean “super-community” size was 119.1 ± 43.1 nodes (ROIs). For intermediate topological scale, *γ* = 1, consensus partitions for all task conditions consisted of three communities with the average size of 89.3 ± 13.2 nodes per community. Consensus partitions detected for the finest topological scale, characterized by the highest structural resolution *γ* = 1.5, consisted of 41.2 communities per partition (average across conditions). This proliferation of observed communities resulted from many singletons, i.e., communities with one node. Each partition consisted of 27-37 singletons, which decreased the average community size to 6.5 ± 16.1 nodes per community. A similar number of communities and average community size characterized condition-independent consensus partitions. For the “super-community” scale, the condition-independent consensus partition consisted of two communities with 141 and 127 nodes per community. The intermediate scale partition was composed of three communities of size 100, 94, and 74 nodes per community. In the finest topological scale, the consensus partition divided the network into 47 communities (38 singletons) with an average community size of 5.7 ± 15.2 nodes per community.

In the coarsest topological scale, the functional network consisted of two major “super-communities”: task-visual and sensory (**Fig. 6**, top panel). The task-visual community was composed of task-related networks (PE signaling, fronto-parietal, memory, and salience networks), default mode network, and visual network. The sensory community consisted of cerebellar, somatomotor, cingulo-opercular, dorsal attention, and auditory networks. Nodes from subcortical and ventral attention networks were divided evenly between two “super-communities.” Intriguingly, in the reward-seeking condition, the visual network was detached from the task-related regions and contributed to the sensory community. Moreover, the positive prediction error partition contained a third smaller subnetwork – ventromedial prefrontal cortex community – unobserved for any other condition (**Fig. 7**, top panel). Four nodes of this community were located within the ventromedial prefrontal cortex – an area widely known for representing value information to drive choice (Hare *et al*., 2009; Rangel *et al*., 2008). The vmPFC community consisted of six regions: three from the default mode network, one from the +PE network, one from the uncertain network, and one from the ventral attention network.

**Fig. 6.**
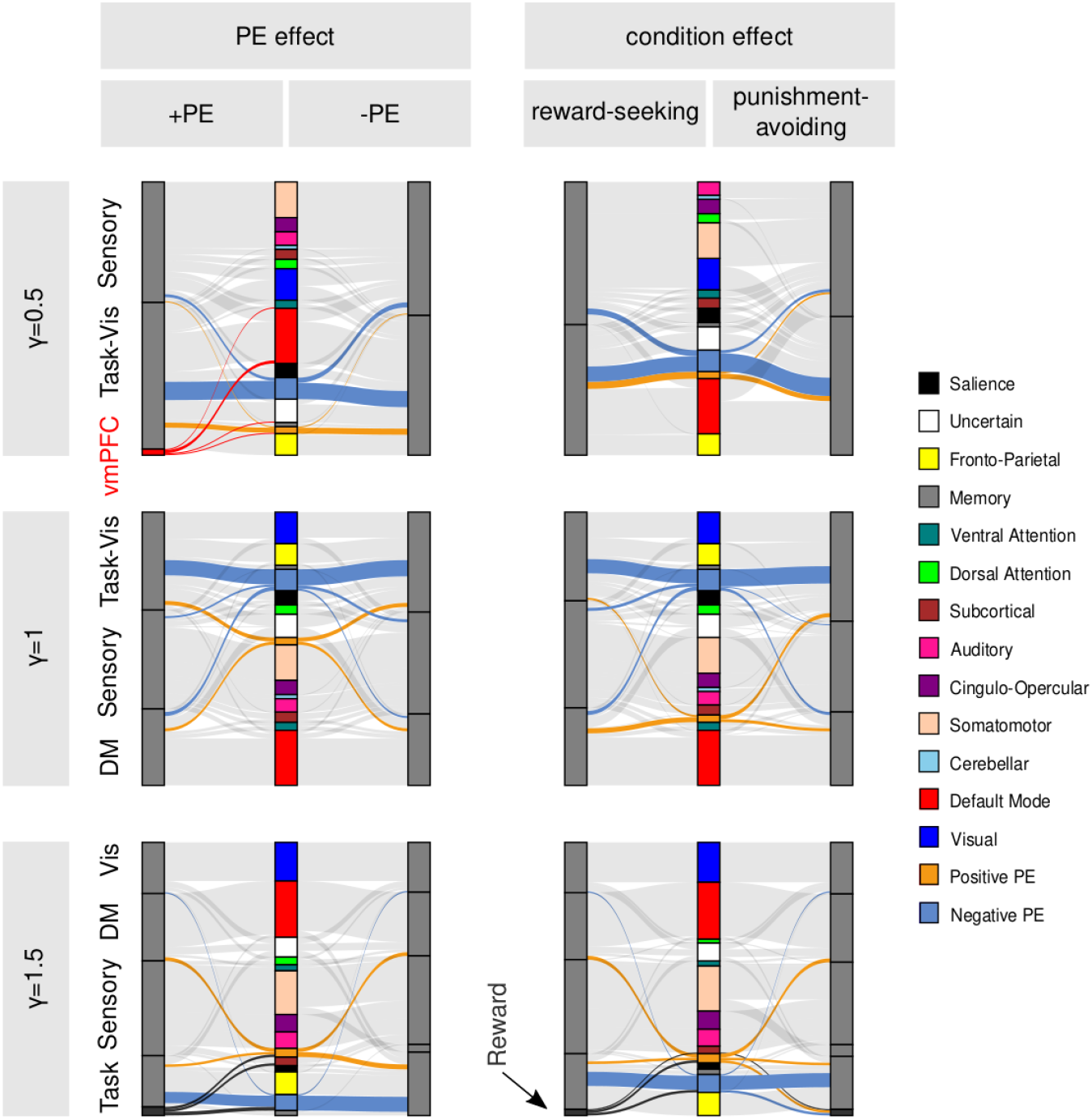
Consensus partitions for different topological scales. Sankey diagrams show composition of consensus partitions with reference to well-known LSNs. First column corresponds to network organization dependent on PE sign, second column shows partition specific to reward-seeking and punishment-avoiding task conditions. Rows correspond to different topological scales characterized by structural resolution parameter γ. Consensus communities: Task-Vis - task-visual; vmPFC - ventromedial prefrontal cortex; DM - default mode; Vis - visual.

**Fig. 7.**
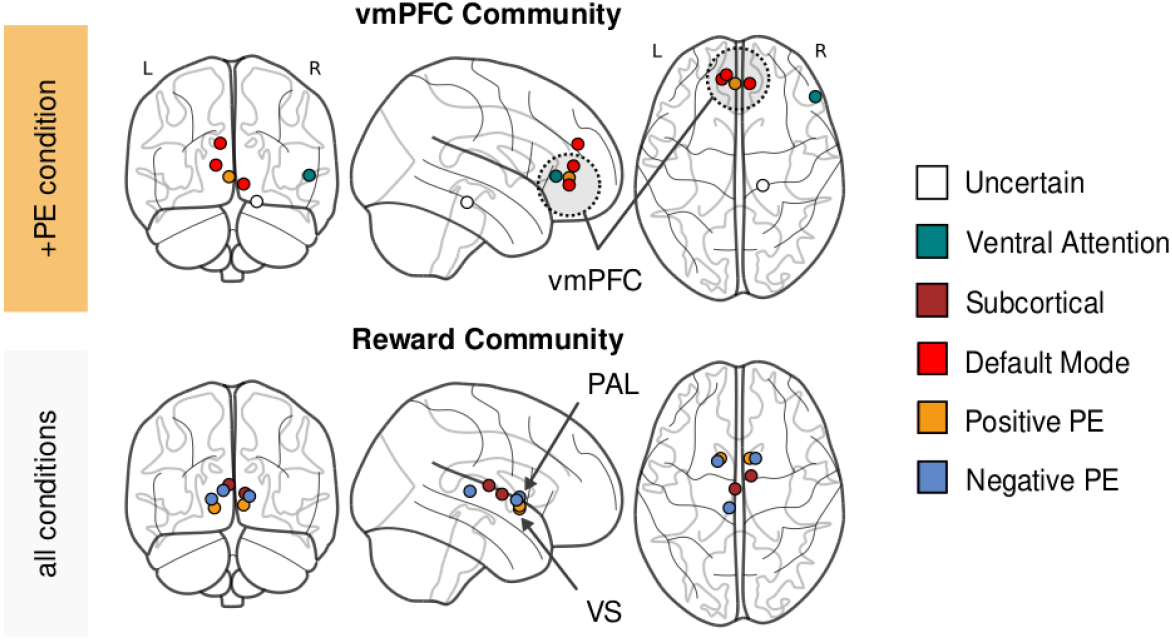
Selected consensus partition communities. (**Top panel**) Ventromedial prefrontal cortex community emerging during processing of positive prediction errors observed for structural resolution γ = 0.5. (**Bottom panel**) Separate reward community observed for condition-invariant consensus partition for structural resolution γ = 1.5. It is comprised of PE signaling regions and subcortical network regions and spatially restricted to the areas around striatum. Regional abbreviations: vmPFC - ventromedial prefrontal cortex; VS - ventral striatum; PAL - pallidum.

In the intermediate topological scale, a pattern of large-scale communities was similar to the one observed in the “super-community” scale (**Fig. 6**, middle panel). One notable difference in network organization between these scales was a separation of the default mode network and part of the +PE signaling network from the task-visual community. The default mode community consisted of most default mode network nodes (85-91%), around half of the +PE signaling, ventral-attention, and uncertain networks nodes, around 10-30% subcortical and 10-20% of −PE signaling and fronto-parietal network nodes. Interestingly, most −PE signaling network nodes remained coupled with the task-visual community. In contrast to the coarsest topological scale, the visual network was consistently coupled with task-related networks for all task conditions.

The analysis of the network partitions for the highest structural resolution, γ = 1.5, showed the fine-grained division of the functional network into many smaller communities. Regardless of experimental condition, the four largest communities—visual, default mode, sensory, and task – formed the backbone of each consensus partition (**Fig. 6**, bottom panel). Similar to the intermediate scale, the default mode community consisted of most default mode network nodes and regions from +PE signaling, ventral attention, and uncertain networks. Task community consisted of regions from fronto-parietal, memory, −PE signaling, +PE signaling and salience network. Interestingly, +PE signaling regions were split between default mode, task, and reward communities, whereas −PE signaling regions resided within the task, reward, and other minor communities. In addition to the backbone communities, a small cerebellar community was consistently discovered for all experimental conditions. Moreover, a stable reward community was a part of the consensus partition for +PE, risk-seeking, and punishment-avoiding conditions. This community was also detected in condition-independent consensus partition representing general network structure during prediction error processing. The reward community consisted of 5-8 regions from PE signaling and subcortical networks (see **Fig. 7**, bottom panel. Specifically, the condition-independent reward community consisted of bilateral ventral striatum and pallidum, right striatum, and two regions within the thalamus. All regions constituting the reward community were spatially bounded to subcortical areas located around the striatum. The existence of a separate reward network was recently suggested by Huckins *et al*. (2019) after examining resting-state connectivity patterns in a large cohort of subjects.

### Large-scale network interactions during prediction error processing

Next we explored condition-specific changes in large-scale networks agreement to test whether modular network architecture fluctuates when subjects switch between processing positive and negative prediction errors. We were curious if such fluctuations can be detected when changing between reward-seeking and punishment-avoiding environments. We hypothesized that changes related to the PE sign should be more pronounced than changes related to the task condition regardless of the structural resolution. We also expected that processing negative prediction errors would increase working memory load leading to an increased between-community agreement and decreased within-community agreement, especially for default-mode, task, and reward systems.

Functional networks in the coarsest topological scale consisted of task-visual and sensory “super-communities” (**Fig. 8**, top panel). Agreement between these “super-communities” was higher during processing −PEs compared with +PEs (D_+PE_ = 0.560, D_−PE_ = 0.579, *t* = −0.63, *p_FDR_* < 0.0001) and higher in reward-seeking compared with the punishment-avoiding condition (D_RS_ = 0.573, D_PA_ = 0.563, *t* = 0.22, *p_FDR_* < 0.0001). Significant change of within-community agreement was observed only for PE sign dimension. Task-visual community was more segregated during processing +PEs as indicated by increased within-community agreement (D_+PE_ = 0.712, D_−PE_ = 0.698, *t* = 0.65, *p_FDR_* = 0.006).

**Fig. 8.**
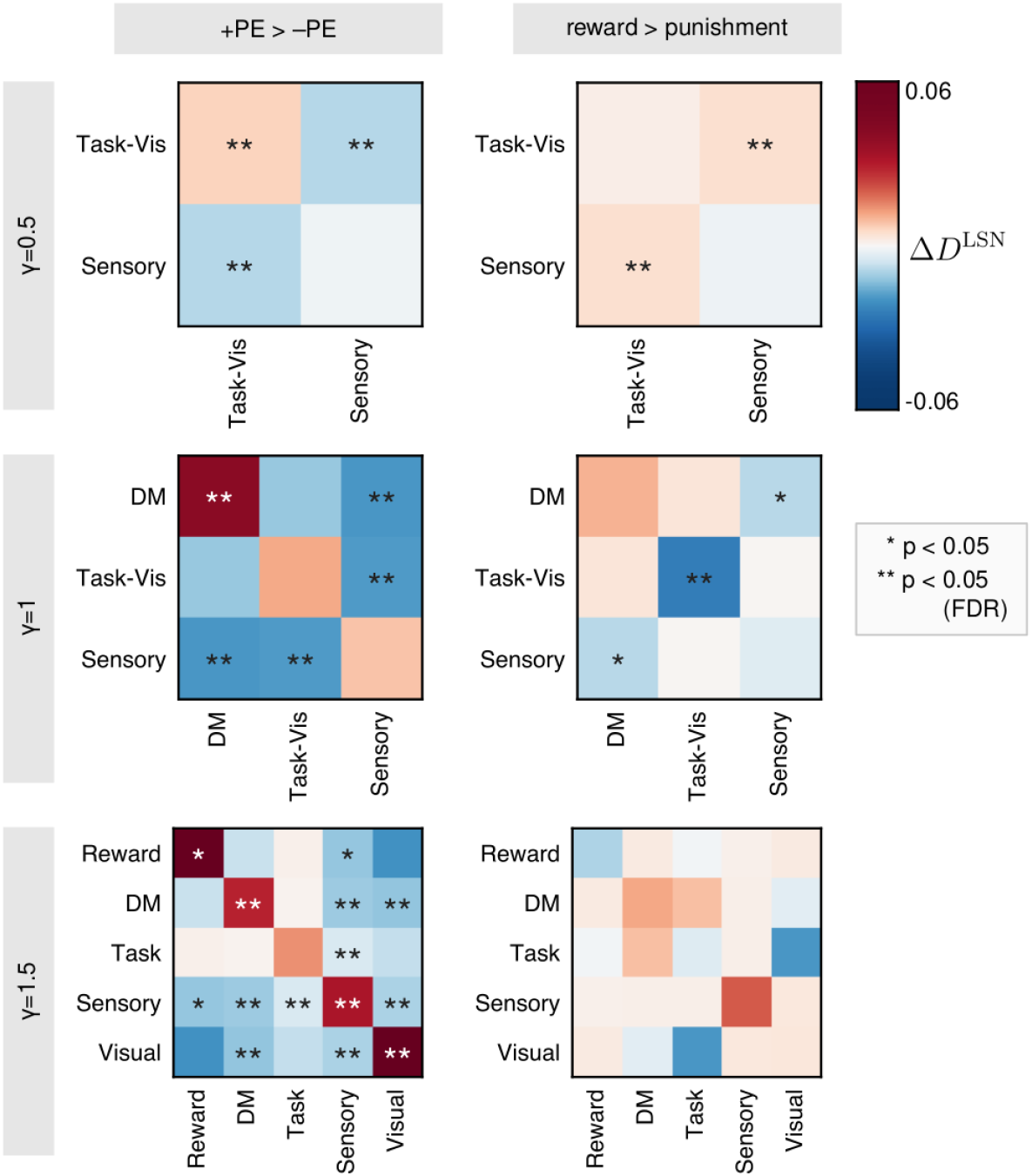
Agreement between large-scale networks. Changes in within and between-community agreement for prediction error sign and task condition. Within-community agreement captures the extent of community segregation whereas between-community agreement quantifies integration between communities. Left column shows the difference between agreement during +PE and −PE processing. Right column corresponds to the changes between reward-seeking and punishment-avoiding conditions. Rows correspond to topological scales. Reference communities come from condition-independent consensus partitions. LSNs abbreviations: Task-Vis - task-visual; DM - default mode.

The intermediate topological scale was characterized by three large-scale communities discovered within the consensus partition: default-mode, task-visual and sensory (**Fig. 8**, middle panel). Three out of six possible changes in the LSN agreement were significant for the PE sign dimension. Agreement within the default mode network increased during processing +PEs compared with −PEs (D_+PE_ = 0.576, D_−PE_ = 0.522, *t* = 3.34, *p_FDR_* < 0.0001). Additionally, the sensory community displayed decreased agreement during +Pes processing with both default-mode (D_+PE_ = 0.183, D_−PE_ = 0.217, *t* = −2.66, *p_FDR_* < 0.0001) and task-visual communities (D_+PE_ = 0.214, D_−PE_ = 0.248, *t* = −3.44, *p_FDR_* < 0.0001). Note that decrease in integration between task-visual and sensory networks was significant for both “super-community” and intermediate scales, whereas increased segregation of task-visual community was confined only to the coarsest topological scale. In contrast to the PE sign effect, only one task-condition change remained significant after correction for multiple comparisons. Agreement within the task-visual community was higher in punishment-avoiding compared with the reward-seeking condition (D_RS_ = 0.514, D_PA_ = 0.555, *t* = −2.19, *p_FDR_* < 0.0001). Interestingly, a similar effect was not observed for the topological scale characterized by γ = 0.5. Increased agreement between default-mode and sensory communities during punishment-avoiding condition was initially significant but did not survive multiple comparison correction (D_RS_ = 0.192, D_PA_ = 0.208, *t* = −1.11, *p_unc_* = 0.04).

The consensus partition for the fine-grained topological scale characterized by γ = 1.5 consisted of nine non-singleton communities (**Fig. 8**, bottom panel). However, only five of these non-singletons comprised more than three network nodes. Since minuscule communities are likely byproducts of a reclustering algorithm without significant biological meaning, they were excluded from further LSN agreement analysis. The remaining communities were labeled as reward, default-mode, task, sensory and visual. None of the 15 possible LSN agreement changes between reward-seeking and punishment-avoiding conditions were significant. Contrary, more than half of PE-sign-related changes turned out to be significant. First, the reward community increased its segregation and decreased its integration with the sensory community with increasing PE. However, any of these changes did not survive multiple comparison correction. Second, Default mode community increased its segregation (D_+PE_ = 0.549, D_−PE_ = 0.503, *t* = 2.04, *p_FDR_* < 0.05) and decreased its integration with sensory (D_+PE_ = 0.061, D_−PE_ = 0.082, *t* = −3.59, *p_FDR_* < 0.0001) and visual communities (D_+PE_ = 0.037, D_−PE_ = 0.061, *t* = −3.31, *p_FDR_* < 0.0001) when switching from negative to positive PEs processing. Third, Task community had a lower agreement with sensory community during +PE processing (D_+PE_ = 0.047, D_−PE_ = 0.056, *t* = −3.54, *p_FDR_*< 0.0001). Third, Sensory community increased within-community agreement with increasing PE (D_+PE_ = 0.483, D_−PE_ = 0.434, *t* = 2.31, *p_FDR_* < 0.0001) and decreased its integration with all remaining communities. Fourth, sensory community increased within-community agreement with increasing PE (D_+PE_ = 0.483, D_−PE_ = 0.434, *t* = 2.31, *p_FDR_* < 0.0001) and decreased its integration with all remaining communities. Fifth, visual community increased segregation during +PE processing (D_+PE_ = 0.530, D_−PE_ = 0.441, *t* = 3.41, *p_FDR_* < 0.001) and decreased integration with default mode and sensory communities.

Almost all changes related to PE sign switching followed a hypothesized pattern of increased within-community agreement and decreased between-community agreement. The only exceptions to that observation were increased agreement during +PE processing between task community and reward/default mode communities for *γ* = 1.5 and decreased task-visual community agreement during +PE processing for γ = 0.5. However, none of these changes reached the significance level, while all significant effects followed the hypothesized pattern. Interestingly, the community separation effect reflected by increased segregation and decreased integration with other communities was more pronounced for sensory, default-mode, and visual communities and less pronounced for task and reward networks.

Prediction error sign effects were more evident than task condition effects for all topological scales, as suggested by more significant changes observed for prediction error sign dimension regardless of the topological scale. This observation was most apparent for the fine-grained topological scale, where none of the task condition effects reached significance in contrast to nine significant effects for the prediction error sign dimension. Moreover, almost all changes related to switching between positive and negative prediction errors were scale-invariant and spanned two or three scales. For example, a decreased agreement between sensory and task/task-visual communities during +PE processing was significant for all three structural resolutions. In contrast, two significant task-condition-related changes were specific to a single structural resolution. Concretely, decreased task-visual community agreement during reward-seeking condition was specific to intermediate scale, and increased agreement between sensory and task-visual communities was significant only for *γ* = 0.5.

## Discussion

Understanding the complex brain mechanisms underlying reinforcement learning is one of the biggest challenges of modern neuroscience. The overarching goal of our study to investigate the functional brain network changes during prediction error processing in the reward-seeking and punishment-avoiding contexts. On the behavioral level, we found that learning speed depends only on the sign of the prediction error and not on the outcome valence. Whole-brain network analysis did not show significant differences in network modularity between processing of negative and prediction errors. Functional networks revealed a multi-scale community structure with a separate reward network emerging at a finer topological scale. The agreement between detected large-scale networks changed between positive and negative prediction error processing. These changes were characterized by decreased within-network segregation and increased between-network integration during negative prediction error processing. This pattern of changes, observed mainly for default mode, sensory and visual networks.

### Opponent system for negative prediction errors processing

From the computational perspective, negative prediction errors are simply negative scalar values located on the same scale as positive prediction errors. This simple observation raises the question: How does the brain represent negative prediction errors? One possible answer to this question takes the form of the dual systems hypothesis stating that negative PEs are signaled by a separate neural system outside of the dopaminergic circuit (Fouragnan et al., 2018). Our network perspective revealed differential community membership profiles of both systems with positive PE system linked with the DMN and negative PE system coupled with the FPN. Behavioral data analysis also supported the duality of PE signaling—the learning rate depended on the sign of prediction error.

Electrophysiological, lesion, and pharmacological studies in animals suggested that the negative part of the prediction error signal is encoded in a set of brain regions outside of the dopaminergic system, most notably in the insula and the amygdala (Hayes *et al*., 2014; Namburi *et al*., 2016). A recent fMRI meta-analysis supported these observations by showing a widespread network of areas signaling negative prediction errors encompassing dorsomedial cingulate cortex, anterior insula, palladium, middle frontal gyrus (Fouragnan *et al*., 2018). A separate community of reward-related regions was recently recognized as a stable large-scale network at rest (Huckins *et al*., 2019). This finding suggests that prediction error processing regions should be functionally coupled. Based on that, we expected that the distinction between regions signaling positive and negative PEs would be reflected by the separate community membership of these networks.

Here, we analyzed community structure on three different topological scales. On the coarsest scale, we found that both prediction error systems were a part of a larger task-visual module. Both intermediate and fine-grained topological scales revealed distinct community membership profiles of positive and negative prediction error processing regions. On the intermediate scale, positive PE regions split evenly between task-visual and default mode modules, whereas negative PE regions retained their association with a task-visual module. The fine-grained scale revealed even more complex structural patterns. We found a separate reward module composed mainly of striatal regions belonging to both prediction error networks. The rest of the prediction error signaling regions were divided between default mode and task modules. The default mode module contained almost half of the positive PE regions and no negative PE regions. On the other hand, the task module consisted of both positive and negative PE regions. These findings partially corroborate the dual system hypothesis (Fouragnan *et al*., 2018) by showing strong associations between the dopaminergic system and DMN and the insular system and FPN. Interestingly, these associations indicate that the opposition between positive and negative PE systems may be related to the antagonism between task-negative DMN and task-positive FPN. Our results show a much more complicated picture of the network community structure in the context of reward-related regions. Most notably, the existence of a separate striatal network comprising both positive and negative PE processing regions suggests a need to integrate information between the opponent systems. We have to note that contrary to the initial hypothesis, we did not find pure modules composed of positive or negative PE processing regions. There are two possible explanations—either some of these regions are irreducibly functionally connected, or the distinction between them exists, but the finer topological scale is needed to uncover them.

Many studies showed that temporal-difference learning models with asymmetric value updates for positive and negative prediction errors are better at explaining learning in humans (Frank *et al*., 2007; Gershman, 2015, 2016). These models were introduced to take into account the observations supporting the dual systems hypothesis (Frank *et al*., 2007). If separate neural circuits signal both types of prediction errors, their influence on choice value should be independent. On the behavioral level, this influence is operationalized as a scalar value—the learning rate. We hypothesized that models with separate learning rates for positive and negative PEs should outperform single learning rate models. In line with our prediction and existing literature, we found moderate evidence in favor of this hypothesis. Moreover, the single most frequent model was the model with PE-dependent learning rates. This finding further suggests the separation between positive and negative PE systems on the behavioral level.

### Brain systems are organized along the prediction error sign axis

The reference effect hypothesis states that values are not absolute but are actively constructed based on our previous experience. For example, we can feel positive emotions in generally negative situations and negative emotions when something good happens. For example, when our car breaks down but repair cost is low, we feel relief, or when our boss praises us, but our colleagues get a raise, we feel envy and frustration. The reference effect hypothesis offers a solution to the ongoing debate on the primary organizational axis of the brain systems. If values were constructed in absolute terms, brain responses would be organized along the outcome valence axis. However, suppose values are relative to the experienced context. In that case, brain responses should only reflect the prediction error sign axis because prediction mirrors the outcome modulated by expectations acting as a reference. To verify the reference effect hypothesis, we investigated whether brain systems are invariant to the outcome valence and prediction error sign. On the behavioral level, we found that learning rates are invariant to the outcome valence but depend on the prediction error sign. On the connectivity level, we found that the prediction error sign effect sizes are much stronger than the outcome valence effect sizes.

The organization of large-scale brain systems constantly changes to meet the demands of the environment (Shine *et al*., 2016). According to the reference effect hypothesis, the functioning of these systems should be invariant to the outcome valence. From the connectivity perspective, this should imply that changes in modular network structure should be predominately associated with switching between positive and negative prediction errors and not between reward-seeking and punishment-avoiding conditions. In agreement with this assumption, we found more significant differences for the change in prediction error sign than the change in outcome valence regardless of the topological scale. Specifically, we found seven significant PE sign effects for the finer topological scale compared with no significant task condition effects. Moreover, switching between different prediction error signs was associated with a topologically stable pattern of large-scale network reorganization. In contrast, outcome valence effects were less consistent and limited to the two coarsest topological scales. Similar to the observed brain activations, these findings support a relaxed version of the reference effect hypothesis—the prediction error axis is a dominant organizational axis for brain systems except for a few subtle valence effects.

### Negative prediction errors elicit brain-wide network reconfiguration

In a reversal learning task, negative prediction errors signal either expected random fluctuations or unexpected changes in reward contingencies. Therefore, negative PE processing may be associated with increased attention and cognitive effort. According to previous studies and theoretical consideration, performing demanding cognitive task requires long-distance connections integrating specialized neural subsystems—so called *global neuronal workspace* (Dehaene *et al*., 1998). Otherwise, the brain can process lower cognitive demands within a set of functionally specialized modules. From the network science perspective, these two effects correspond to decreased modularity, increased within-community integration, and decreased between-community segregation. Several studies reported these changes in functional brain networks during cognition (Vatansever *et al*., 2015, Finc *et al*., 2017; Shine *et al*., 2016). Similar to previous studies, we expected to observe decreased whole-brain modularity, within-community segregation and increased between-community integration when switching from positive to negative PE processing. In line with these expectations, we observed decreased segregation of default mode, sensory and visual modules, and increased integration between default mode and sensory/visual modules during negative PE processing. However, overall network modularity was stable across prediction error signs and task conditions.

Although the whole-brain modularity values were lower for negative PEs in all three topological scales, none of these differences reached a significance level. There are two possible explanations for that observation. First, in the case of probabilistic learning, cognitive effort differences between positive and negative PEs could be much lower than similar differences during working memory or attention tasks. Subtle differences in cognitive effort would lead to a smaller modularity breakdown that cannot be detected with the sample size used in our study. This explanation is consistent with the observation of lower modularity for negative PEs in all topological scales and moderately small p-value (*p* = 0.18) for PE effect in the “super-community” topological scale. Second, the reorganization effect may be specific to changes in the composition of network communities without altering the macroscopic level of global modularity (Sporns, 2014). Similar to our results, a recent study on number comparison reported significant alterations of network community structure without changes in whole-brain network modularity (Conrad *et al*., 2020).

In the coarsest topological scale, the default mode network was a part of the task-visual module. This module decreased its segregation and increased integration with the sensory module during negative PE processing. In intermediate and finer topological scales, DMN formed a separate module. Similar to the task-visual module, this module decreased segregation and increased integration with sensory and visual modules. Recent work demonstrated that the DMN may play an crucial role in the formation of an integrated workspace (Finc *et al*., 2017). In the context of reinforcement learning, DMN contains a part of the prediction error signaling network, most notably the ventromedial prefrontal cortex commonly associated with value representation (Frank & Claus, 2006; Rangel *et al*., 2008). Our findings suggest that DMN reorganization patterns may reflect (1) global workspace formation and (2) enhanced communication between the valuation circuit and other brain systems following negative PE processing.

The sensory network contains the motor cortex responsible for the movement execution following the subject’s decision. Some studies suggest that motor regions may implement a winner-take-all mechanism integrating values of different stimuli (Cisek & Kalaska, 2005). Moreover, Horga *et al*. (2015) demonstrated the importance of sensorimotor connectivity for reinforcement learning during gradual learning in a virtual maze. Interestingly, our results show that the sensory network increases its agreement during negative PE processing with all other systems regardless of the scale. This suggests a potential need for information integration between the choice and valuation circuit and other large-scale networks.

We observed the separate reward network only for finer topological scale characterized by resolution parameter *γ* = 1.5. The reward network consisted of a few striatal regions signaling both types of prediction errors. Consistently with the reconfiguration pattern of the global workspace formation, the reward network decreased its segregation and increased its integration with the sensory network during negative PE processing. Decreased segregation of striatal regions may reflect functional decoupling of positive and negative PE processing regions during a more effortful type of prediction error processing. This decoupling possibly allows reducing interference between antagonistic areas when more cognitive resources are required. We have to note that reward system effects did not survive multiple comparison corrections and need to be interpreted with caution. A smaller effect size for the reward network may be related to a smaller community size of only seven nodes.

Agreement analysis revealed only two changes in large-scale network integration and segregation for the outcome valence axis. Both changes were related to the task-visual module and were observed for the two coarser topological scales (*γ* = 0.5 and *γ* = 1). This module increased its integration with the sensory module and decreased its segregation during the reward-seeking condition. One possible explanation of this effect is the influence of systematic differences in stimulus presentation during the outcome phase described in the previous section. In the activation analysis, we observed differences in PE processing in a large portion of the primary and secondary visual cortex, a part of the task-visual module. It is also important to note that despite statistical significance, both effects were scale-specific and were not present for other topological resolutions, unlike PE-related changes.

### Ventromedial prefrontal regions form separate network during positive prediction error processing

The ventromedial prefrontal cortex is one of the main functional hubs of the default mode network (Andrews-Hanna *et al*., 2014). From the reinforcement learning perspective, vmPFC signals positive prediction errors (Daw *et al*., 2011; Van den Bos *et al*., 2012) and represents a subjective value of various stimuli (Levy & Glimcher, 2012; Roy *et al*., 2012). Using consensus partitioning, we found that regions spanning the vmPFC area formed a separate network community during positive prediction error processing.

Surprisingly, this community was present only for the coarsest topological scale (*γ* = 1.5), which inherently favors larger “super-communities.” This may suggest remarkably strong connections among vmPFC nodes during positive PE processing. It has been previously suggested that value representation in vmPFC is updated by striatal prediction errors through strong frontostriatal connections (Frank & Claus, 2006). Connectivity studies supported this claim by showing increased functional coupling between ventral striatum and vmPFC during feedback processing (Camara *et al*., 2009; Münt*e et al*., 2008). Moreover, Van den Bos *et al*. (2012) showed that connection strength between these two regions is enhanced during positive feedback processing. This value update mechanism offers a possible explanation of our finding – the dopaminergic system may communicate positive prediction errors with vmPFC through frontostriatal connections evoking strong and coherent neural activity in different parts of vmPFC. Strong coherent activity results in strong functional coupling, which results in a separate vmPFC community observed during positive PE processing. The other complementary mechanism for community formation is decreased connectivity with non-community members. From the PE processing perspective, this would reflect the need for the valuation circuit in vmPFC to reduce noise from other brain systems during value updating. Only one or both of the suggested mechanisms may be responsible for the vmPFC community formation. Therefore more studies and fine-grained network structure analysis would be needed to understand this phenomenon better. Since this result was unexpected, it should be treated as exploratory and interpreted with caution.

## Conclusions

Multiple neural systems are engaged to signal prediction errors enabling us to learn through trial-and-error. The results of our study show that these systems are organized along the prediction error sign axis, reflecting the distinction between positive and negative prediction errors. Moreover, the activity and connectivity of these systems are generally invariant to the outcome valence, supporting the idea that the brain adjusts its reference point when the environment is predominantly rewarding or punishing. Our results also demonstrate a complex pattern of network interactions following switching between positive and negative prediction errors. These interactions form the pattern observed in studies on cognitive load, suggesting that negative prediction errors require more cognitive resources as they are usually related to the conflict. Intriguingly, we also found some unexpected network community structures. Ventromedial prefrontal regions formed a small community observed along two large “super-communities,” suggesting increased integrity of the valuation system and decreased communication with the rest of the brain during positive prediction error processing. Striatal areas composed of both regions signaling positive and negative prediction errors formed a stable reward network observed for higher topological resolutions supporting the finding of a separate reward network observed at rest. Our findings provide a thorough description of neural correlates of prediction errors from three different perspectives. They support and extend some existing observations and shed new light on the network mechanism behind reinforcement learning. Some of our results raise new fascinating questions and open avenues for future research.

## Methods

### Participants

Thirty-two healthy volunteers (14 female; mean age: 20.9 ± 2.24; age range: 18-28) were recruited from the local community through social networks and word-of-mouth. All participants were right-handed, had a normal or corrected-to-normal vision, and did not suffer from neurological or psychiatric disorders at the time of examination or in the past. Informed consent was obtained in writing from each participant, and ethical approval for the study was obtained from the Ethics Committee of the Nicolaus Copernicus University Ludwik Rydygier Collegium Medicum in Bydgoszcz, Poland.

### Experimental procedures

Probabilistic reversal learning (PRL) task was selected to examine neural correlates of prediction errors (Cools *et al*., 2002). The PRL task consisted of two conditions (1) reward-seeking and (2) punishment-avoiding. In each of these variants, winning and losing are opposed to neutral outcomes, enabling disentangling between often confused dimensions: outcome valence and prediction error sign (Palminteri & Pessiglione, 2017). Both task conditions were performed inside the fMRI scanner. Participants were instructed to repeatedly choose between yellow and blue boxes to collect as many points as possible in the reward-seeking condition or lose as few points as possible in the punishment-avoiding condition (**Fig. 1A**). One of the boxes had the probability of being correct (rewarding or non-punishing depending on the condition) *p* = 0.8 and the other one *p* = 0.2. These probabilities were unknown to the subjects and had to be learned from experience. The reward/punishment contingency changed four times throughout each task condition (**Fig. 1C**). Each box had associated reward/punishment magnitude, randomly selected at the beginning of each trial. Magnitudes for both boxes were integers summing up to 50, with the difference between them not exceeding 40. They were represented as white numbers on the boxes, indicating possible gain in the reward-seeking condition or loss in the punishment-avoiding condition. Successful performance in the PRL task requires the decision-maker to correctly estimate correct choice probabilities from experience and integrate them with reward/punishment magnitudes to choose an option with a higher expected value.

Each task condition was associated with the separate fMRI run and consisted of 110 trials. Each trial began with the decision phase indicated by the question mark appearing within the fixation circle (**Fig. 1B**). During the decision phase a subject had 1.5 s to choose one of the boxes by pressing a button on the response grip with either left or right thumb. The decision phase was followed by a variable inter-stimulus-interval (ISI; 3-7 s, jittered), after which an outcome was presented for 1.5 s. During the outcome phase, the fixation circle was colored according to the rewarded or punished box, and the number within the circle represented the number of gained or lost points. The outcome phase was followed by a variable inter-trial-interval (ITI; 3-7 s, jittered).

The gray account bar on the bottom of the screen represented points that a subject gathered in the reward-seeking condition or the remaining points in the punishment-avoiding condition. In the reward-seeking condition, subjects were informed that if they fill half of the bar or the entire bar, they will receive 10 PLN (≈ 2.5 USD) or 20 PLN (≈ 5 USD), respectively. Similarly, in the punishment-avoiding condition, they were informed that they would receive 20 PLN if left with more than half of the bar, 10 PLN if left with less than half of the bar, and no money if they lose all of their points. To maintain a constant motivation throughout the task, incentives thresholds were set such that all participants acquired 10 PLN from either task.

Heterogeneity in the prior expectations regarding the task structure may lead to heterogeneity in behavior even in simple tasks leading to inaccurate behavioral modeling (Shteingart & Loewenstein, 2014). Therefore, participants were explicitly instructed that one of two boxes (without telling which) will be more frequently rewarded in the reward-seeking condition or punished in the punishment-avoiding condition and that this contingency may reverse several times throughout the task. Before the MRI scan, subjects practiced both task conditions on the lab computer. During the first phase of the practice, participants were provided with feedback indicating which box is more frequently correct to ensure that they grasp the correct model of the task environment.

PsychoPy software (v. 1.90.1; www.psychopy.org; Peirce (2007)) was used for task presentation on the MRI compatible NNL goggles (NordicNeuroLab, Bergen, Norway). Behavioral responses were collected using MRI-compatible NNL response grips (NordicNeuroLab, Bergen, Norway), which were held in both hands. Each condition lasted approximately 24 min. The order of task conditions and the colors for the left and right boxes (yellow and blue) were counterbalanced across subjects.

### Behavioral model space

Reinforcement learning models can quantitatively account for learning by trial-and-error (Montague, 1996). Precisely, models based on the idea of the temporal-difference learning postulate that stimulus values are updated proportionally to the prediction error weighted by adjustable learning rate (Wagner & Rescorla, 1972). Because of the anticorrelated design of correct choice probabilities during the PRL task, simultaneous stimulus value updates for the chosen and non-chosen options were assumed (O’Doherty *et al*., 2007). Following previous studies, models with separate learning rates for positive and negative PEs were considered to account for possible risk-sensitivity effects. These models would be referred to as prediction-error dependent (PD) models (Niv *et al*., 2012; Reiter *et al*., 2016; Van den Bos *et al*., 2012). To capture possible valence effects, e.g., loss aversion, models with separate learning rates for reward-seeking and punishment-avoiding conditions were also included. These models would be referred to as condition-dependent models (CD). To fully explore model space, all possible combinations of models were considered, which resulted in four different models:

1. PICI model with single learning rate independent of the sign of prediction error and outcome valence,
2. PICD model with two separate learning rates for reward-seeking and punishment-avoiding conditions,
3. PDCI model with two separate learning rates for positive and negative PEs,
4. PDCD model with four separate learning rates for prediction error signs and task conditions.

All models assumed that prediction error at each trial t is computed as the difference between experienced outcome, r t, and the expected probability that chosen option will be correct (rewarding / not-punishing), *p_t_^c^*

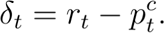

Outcomes could be either *r_t_* = 1 when a stimulus is rewarded / not punished or *r_t_* = 0 when stimulus is punished / not rewarded at trial t. Note that randomly drawn reward magnitudes did not follow any structured rules. Hence, participants learned and estimated only correct choice probabilities and not action values. In each trial, probability estimates for both chosen, *p_t_^c,^* and unchosen stimulus, *p_t_^u^* were simultaneously updated according to a standard Rescorla-Wagner rule:

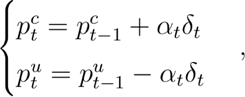

where *α_t_* ∈ [0, 1] is the learning rate used in trial *t*. Note that learning rate values close to zero result in minor updates and slow learning, whereas values close to one result in probability matching behavior. In the simplest PICI model, a single learning rate, *α_t_^PICD^*, was used to update probability estimates regardless of PE sign and task condition. The rest of the models assumed separate learning rates depending on task condition, prediction error sign, or both:

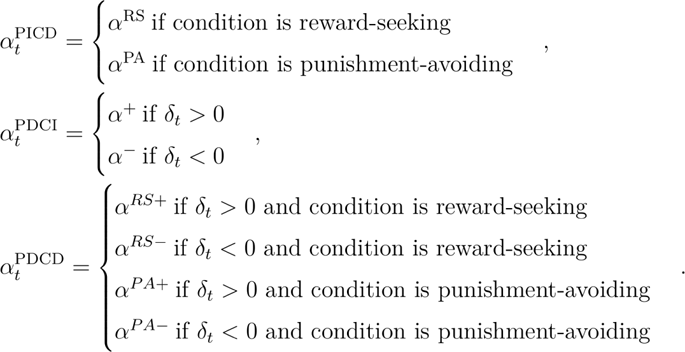

Action selection was modeled based on reward/punishment magnitudes and continuously updated probability estimates. Utility of both options was assumed as Pascalian expected value, i.e., a product of the expected probability that the box is rewarded/punished, ρ_*t*_, and reward/punishment magnitude for that box, *x_t_*:

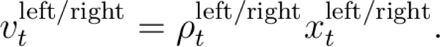

Note that in the case of punishment-avoiding condition expected probability that stimulus leads to punishment equals one minus the expected probability that it is a correct choice:

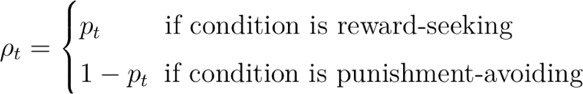

Finally, the choice probability was derived by coupling expected values with the softmax policy rule (Luce, 1957):

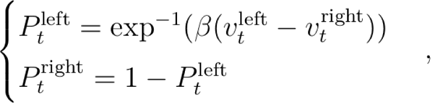

where precision (or “inverse-temperature”) parameter *β* ∈ [0, ∞) reflects choice stochasticity by controlling sensitivity of the choice probability to differences in expected value between the two stimuli. Choice probability values served as likelihood functions generating choices in the Bayesian modeling framework.

### Behavioral performance

Successful performance in the PRL task requires subjects to take into account changing reward or punishment probabilities and magnitudes associated with each choice. As in any version of the PRL task, probabilities had to be learned from experience by trial-and-error. An individual subject’s performance accuracy was calculated to establish if subjects succeeded in learning probabilistic associations. Accuracy was defined as the proportion of choices that led to rewarding outcomes in reward-seeking condition or non-punishing outcomes in punishment-avoiding condition. Since the study examined healthy subjects without valence-dependent learning deficits, we hypothesized similar performance levels for both task conditions.

### Bayesian modeling

Bayesian hierarchical latent-mixture (HLM) model was used to perform model selection and parameter estimation. This type of Bayesian model contains parameters representing behavioral parameters like learning rates or precision and parameters reflecting the belief about which competing model generates the observed data. The hierarchical structure of the model assumes behavioral parameters of individuals coming from group-level distributions. The latent-mixture structure assumes that observed behavior can arise as a combination of computations from different cognitive processes. The HLM model was estimated with PICI, PICD, PDCI, and PDCD submodels as competing submodels for model comparison (**Fig. S1**). Then a hierarchical model with a winning submodel as a generative model for behavioral responses was estimated for parameter recovery purposes.

Weakly informative hyperpriors were set to model the distribution of the learning rates and precision across individuals. Following previous studies, learning rates were assumed to come from a beta distribution (Gershman, 2016). Beta distribution is defined for the continuous interval α ∈ [0, 1] covering entire range of possible learning rate values:

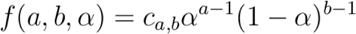

where a and b are two non-negative shape parameters. These were set to come from the uniform hyperprior distribution *a* ∼ Uniform(1, 10) and *b* ∼ Uniform(1, 10). Note that both shape parameters were assumed greater than 1, since for *a* < 1 and *b* < 1 beta distribution takes a behaviorally implausible U-shape. Precision parameter *β* was independently included for each submodel to provide additional flexibility in model parameter space and avoid potential dependence between competing models. Generative distribution for precision parameter *β* was represented as a lognormal distribution. Hyperpriors for lognormal parameters distributions were set to *µ* ∼ Uniform(−2.3, 3.4) for lognormal group mean and σ ∼ Uniform(0.01, 1.6) for lognormal standard deviation. These weakly informative hyperpriors were used in previous studies and derived from the constraints to confine group mean for the precision parameter within 0.1 to ∼ 30 interval (Meder *et al*., 2019; Nilsson *et al*., 2011). The latent-mixture part of the model assumed a subject-specific model indicator variable to represent a latent mixture of four competing submodels. This assumption allowed to model each subject response pattern independently by using a different mixture of competing submodels. Model indicator variable, *z_i_*, was modeled with uniformly distributed flat prior representing lack of expectations toward which behavioral model best explains subjects’ response patterns.

The *Markov Chain Monte Carlo* (MCMC) sampling was performed using JAGS software (v4.3.0; http://mcmc-jags.sourceforge.net/) called from MATLAB R2017a via matjags.m script (v1.3.3; https://github.com/msteyvers/matjags). A sampling procedure with 2000 burn-in samples, four chains, and 15,000 samples per chain was used to estimate the hierarchical latent-mixture model (see **Supplementary Information**). The single best submodel was selected by calculating estimated model frequencies and protected exceedance probabilities (Rigoux *et al*., 2014) for the model frequency being highest among other submodels. These calculations were performed with the Variational Bayesian Analysis toolbox (v1.9.2; https://github.com/MBB-team/VBA-toolbox), which implements Bayesian model selection for group studies. Model parameters for the winning model were fitted by estimating a reduced hierarchical latent-mixture model.

The HLM model with four competing submodels was evaluated to select the winning submodel. A sampling of the probability distribution of the model indicator variable z i resulted in posterior distribution for each subject reflecting posterior probability for each competing submodel.

### Data acquisition

Brain imaging data were collected using a GE Discovery MR750 3 Tesla fMRI scanner (General Electric Healthcare) with a standard 8-channel head coil. Anatomical images were obtained using a three-dimensional high resolution T1-weighted (T1w) gradient-echo (FSPGR BRAVO) sequence (TR = 8.2 s, TE = 3.2 ms, FOV = 256 mm, flip angle = 12 degrees, matrix size = 256 × 256, voxel size = 1 × 1 × 1 mm, 206 axial oblique slices). Functional images were obtained using a T2*-weighted gradient-echo, echo-planar imaging (EPI) sequence sensitive to BOLD contrast (TR = 2,000 ms, TE = 30 ms, FOV = 192 mm, flip angle = 90 degrees, matrix size = 64 × 64, voxel size 3 × 3 × 3 mm, 0.5 mm gap). Two runs of probabilistic reversal learning in the reward-seeking and punishment-avoiding conditions were acquired (24 min 20 s; 730 volumes each). Forty-two axial oblique, interleaved slices were scanned for each functional run, and five dummy scans (10 s) were collected at the beginning of each run to stabilize magnetization.

### Data processing

The raw DICOM data were converted to NifTI format, structured according to the Brain Imaging Data Structure (BIDS) standard, and validated using BIDS Validator (https://bids-standard.github.io/bids-validator/) (Gorgolewski *et al*., 2016; Yarkoni et al., 2019). The preprocessing was performed using fMRIPrep version 1.4.1 (Esteban *et al*., 2019; Gorgolewski *et al*., 2011) (see **Supplementary Information** for the full description of pipeline).

### Brain parcellation

To extract brain signals we used parcellation introduced by Power *et al*. (2011) that consists of 264 regions of interest defined based on a meta-analysis of activation patterns from multiple studies. Each ROI is modeled as a 5mm sphere around the specific MNI coordinates. Power’s parcellation additionally includes the division of the 264 ROIs into 13 large-scale networks: auditory, cerebellar, somatomotor, cingulo-opercular, default mode, memory, ventral attention, dorsal attention, fronto-parietal, salience, subcortical, uncertain and visual. This division allows examining the dynamic interplay between LSNs in various experimental conditions. However, Power’s parcellation does not contain separate prediction-error-related networks; therefore, it cannot be directly used to study how the reward system interacts with the rest of the brain during reinforcement learning. Here, the Power’s ROIs were combined with regions from the recent meta-analysis on neural representations of prediction error valence (Fouragnan *et al*., 2018) (see **Supplementary Information** for details).

### Beta-series correlation

Task-related functional connectivity was estimated using a beta-series correlation (BSC) method introduced by Rissman *et al*. (2004). The BSC method quantifies event-to-event fluctuations in the activity of different brain areas. These fluctuations allow estimating statistical dependency between regional activations for different event types. The “single-trial-versus-other-trial” approach introduced by Mumford *et al*. (2012) was used to compute activation beta maps. This approach assumes creating a separate GLM for each trial in which the trial of interest is modeled as a separate regressor, and all other trials are modeled as a single nuisance regressor. Here we used the FirstLevelModel class from the Nistats package and a custom code to calculate beta-series correlation. For each trial, a separate GLM was created. The GLM consisted of (1) one regressor modeling a trial of interest as an event of duration 0 s (typically used to model very short cognitive processes) time-locked to the onset of the outcome phase, convolved with a standard SPM hemodynamic response function (HRF), (2) two regressors modeling trials with positive and negative prediction error as events of duration 0s time-locked to the onset of the outcome phase, convolved with an HRF, (3) six regressors modeling the events of no interest: decision onset, decision onset modulated with expected probability for side for being correct, decision onset for missed trials, left button press, right button press, and decision offset for missed trials, (4) twenty-four head motion parameter regressors modeling motion-related noise, (5) cosine functions modeling low-frequency scanner drift. During the GLM timerseries were high-pass filtered with the cut-off frequency 128 Hz (see **Supplementary Information** for details).

The output of this procedure consisted of a three-dimensional beta map for each trial, indicating the voxel-wise estimate of neuronal activity evoked during this trial. After GLM estimation, beta values for each trial were averaged in each of 268 ROIs producing 268 beta-series per subject and task condition. Beta-series were then z-transformed and separated by the sign of the prediction error of each trial. This procedure gave four beta-series for each participant: +PE trials in reward-seeking condition, −PE trials in reward-seeking condition, +PE trials in punishment-avoiding condition, and −PE trials in punishment-avoiding condition.

Beta-series were then correlated using Pearson’s correlation coefficient, generating a *N ROI × N ROI* symmetrical correlation matrix for each subject, task condition, and trial-wise prediction error sign. Correlation values were then converted to z-scores using Fisher z-transform. Transformed matrices represented the final subject-level estimate of functional connectivity pattern evoked by the specific task condition and type of trial (see **Supplementary Information** for details).

### Multi-scale modularity

To calculate community structure properties across topological scales, the structural resolution parameter space was sampled by varying *γ* from 0.05 to 3 with a step of 0.05. For each value of γ, the Louvain modularity maximization algorithm was used to quantify optimal network structure (Blondel *et al*., 2008). Negative connections were separately included in the modularity function and treated asymmetrically as suggested in Rubinov and Sporns (2011):

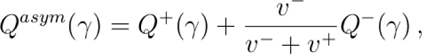

where *Q* ± (*γ*) are modularity values calculated separately for positive and negative connections, and *v* ± = *ij A* ± *ij* are total connection strengths, i.e., costs for positive and negative connections. For each functional network, the Louvain algorithm was run 100 times, and the partition with the highest value of *Q asym* (γ) was kept for further analysis. Several metrics were computed to inspect different topological scales (**Fig. S6**): (1) a mean number of communities, singletons, and non-singletons (communities with more than one node), (2) mean community size, (3) mean similarity between data-driven partitions, and (4) mean similarity between reference partition and data-driven partitions. Partition similarity was computed as the normalized mutual information between community partition vectors (Meilă, 2007). A sparse sampling of γ range was employed, enabling to focus on three distinct topological scales: *γ* = 0.5 representing the largest scale of few “super-communities” consisting of multiple LSNs, *γ* = 1 representing most frequently studied intermediate scale, and *γ* = 1.5 representing finer scale approximately resembling well-known division into LSNs (see **Supplementary Information** for details).

To assess subject-level modularity and community structure for both task conditions and trial-wise prediction errors signs, we employed a similar procedure to the one used during structural resolution parameter analysis. For each value of *γ* ∈ {0.5, 1, 1.5} and each functional network, the Louvain modularity maximization algorithm was executed 1000 times, and the output of the run with the highest modularity was saved for further analysis. The output consisted of weighted modularity with the asymmetric treatment of negative weights, *Q asym* (called *Q* in subsequent analysis), and a *N ROI* × 1 community affiliation vector, *M*, whose i-th element indexed community affiliation of node i.

### Consensus partitioning

A consensus partition represents a modular structure of a functional brain network during the specific experimental condition. Group-level community structure represented by consensus partition can be calculated from subject-level community affiliation vectors. Consensus partitions effectively decrease data dimensionality, allowing a qualitative description of the network reconfiguration. There are two popular approaches for defining a consensus partition: (1) selecting a partition with the highest similarity to the other partitions (Doron *et al*., 2012), and (2) reclustering an association matrix that stores the information about pairwise nodal co-occurrence within the same community (Bassett *et al*., 2013; Betzel *et al*., 2017; Lancichinetti & Fortunato, 2012). Here, the latter approach based on the concept of agreement (or module allegiance) was used. The agreement, *D_ij_*, is a measure defined for the pair of nodes, *i* and *j*, and characterizes the extent to which these nodes belong to the same community (Bassett *et al*., 2015; Bertolero *et al*., 2015):

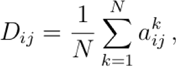

where *N* is the number of partitions, and a *k_ij_* equals 1 if nodes i and *j* belong to the same community, and 0 otherwise. The elements of the agreement matrix, *D_ij_*, reflect the probability that nodes *i* and *j* are found within the same community for randomly selected partition given a set of partitions. Although information represented by the agreement matrix is not independent of the information contained in the connectivity matrix, an agreement can provide a complementary characteristic of nodal associations. For example, two nodes can share a weak direct connection but many strong indirect connections. In this case, there is a high chance that they will be placed within the same community exhibiting significant agreement and low connectivity at the same time. Furthermore, Bassett *et al*. (2015) has shown that agreement reduces the noise in connectivity matrices and is more sensitive to differences in network organization compared with information contained in connectivity values alone.

Representative modular structure for task conditions and prediction error signs was calculated in four steps (**Fig. 3**). First, subject-level community affiliation vectors were aggregated into four groups: reward-seeking, punishment-avoiding, positive PE, and negative PE. Each group consisted of two vectors per subject, i.e., 58 in total. For example, a reward-seeking group consisted of partitions for both +PE and −PE networks in reward-seeking condition. Similarly, a positive PE group consisted of +PE networks pooled over reward-seeking and punishment-avoiding conditions. This pooling procedure allowed us to examine representative community structure independently for both experimental dimensions. Additionally, the fifth group consisting of partitions from all task conditions was created. It consisted of four vectors per subject, i.e., 116 in total. This group was included to represent a modular network structure during prediction errors processing that is stable across task conditions. Second, an agreement matrix was computed for each group. Third, agreement matrices were thresholded to remove weak associations, i.e., all agreement values below *τ* = 0.5 were set to zero. Finally, thresholded agreement matrices were iteratively reclustered with the Louvain algorithm until convergence, and the output affiliation vectors were considered consensus partitions. This procedure yielded four condition-specific and one condition-independent community vector. The consensus partitioning procedure was independently repeated for three values of structural resolution parameter *γ*. Agreement matrices and consensus partitions were calculated using `agreement` and `consensus_und` functions from bctpy package (version 0.5.2) based on the Brain Connectivity Toolbox (Rubinov & Sporns, 2010).

### Community-level agreement

To investigate condition-specific changes in the community-level agreement, individual community affiliation vectors were used to calculate individual node-level agreement matrices (**Fig. 4**). An individual node-level agreement is an *N ROI × N ROI* matrix with entries *D_ij_* ∈ {0, 1} indicating whether a pair of nodes belongs to the same data-driven community. These node-level matrices were then block-averaged according to the network division into LSNs in condition-independent consensus partition to create individual LSN agreement matrices:

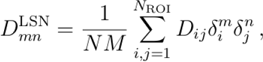

where *N* and *M* are the sizes of communities *n* and *m*, and δ is a community indicator equal to 1 if node *i* belongs to community m and 0 otherwise. Then, for LSN each community-level agreement value, *D_mn_*, a two-sided paired t-test was conducted for task condition and prediction error sign effects. The null hypothesis states that the degree of LSN agreement is the same in reward-seeking and punishment-avoiding conditions for a task condition effect. For a prediction error sign effect, the statistical test aims to answer if there is a significant change of LSN agreement when switching between signaling positive and negative prediction errors.

To determine the significance of the computed t-statistics, a Monte Carlo permutation procedure was employed (Conrad *et al*., 2020). Permutation-based procedures represent an alternative approach to significance testing that is especially useful in multidimensional neuroimaging data (Nichols & Holmes, 2002). The total number of 10,000 iterations was performed to obtain a reliable estimate of the null distribution for computed t-statistics. For each iteration, the subject-level community allegiance vectors, M, reflecting modular network structure, were randomly shuffled to represent a random topology unrelated to the observed functional connections while preserving the number and size of the original communities. These vectors were then subjected to the same procedure employed for the actual dataset consisting of agreement calculation, block-averaging, and statistical testing. Each iteration outputted two null t-statistic matrices of size *N LSN × N LSN* for both effects of interest. A set of 10,000 of these matrices formed a null distribution of the t-statistic. P-values were then calculated based on the null distribution percentile corresponding to the true t-statistic. An effect was considered significant if the true t-statistic was lower than 2.5th or higher than 97.5th percentile of the null distribution. A two-sided approach was used to account for both increases and decreases in a community-level agreement between conditions. Permutation-based p-values were then corrected for multiple comparisons using the FDR method (Benjamini & Hochberg, 1995).

## Data and code availability

All code used for neuroimaging and behavioral data processing and statistical data analyses are publicly available at https://github.com/kbonna/decidenet. The raw fMRI data are available from the corresponding author on request.

## Acknowledments

The study was supported by the National Science Centre, Poland (2020/36/T/HS6/00104). K.B, K.F. & W.D. were supported by *The Excellence Initiative—Research University* awarded to Nicolaus Copernicus University.

## 1. Behavioral modeling

### 1.1. Bayesian modeling

Bayesian modeling provides a strict and flexible way of relating formal cognitive models to behavioral data (Lee & Wagenmakers, 2014). Bayesian modeling aims to estimate a set of parameters, *θ*, which can represent hidden parameters of behavioral models or variables used for model comparison. In Bayesian statistics, parameters are represented as probability distributions instead of single numbers, which preserves the information about their uncertainty. The general principle of Bayesian analysis is to use collected data, *X*, to update the prior beliefs about parameters to posterior beliefs represented as new probability distributions over the parameters in question. The “updating process” is captured in a Bayes formula:

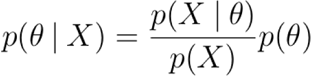

where *p*(*θ* | *X*) is a posterior distribution of *θ* given the data *X*, *p*(*X* | *θ*) is a probability of observing data *X* given prior beliefs about *θ*, *p*(*θ*) is a prior probability distribution, and *p*(*X*) is a normalizing constant. Posterior distribution *p*(*θ* | *X*) represents updated belief about parameters and takes into account both prior beliefs and collected data *X*.

Weakly informative hyperpriors were set to model the distribution of the learning rates and precision across individuals. Following previous studies, learning rates were assumed to come from a beta distribution (Gershman, 2016). Beta distribution is defined for the continuous interval *α* ∈ *[0, 1]* covering the entire range of possible learning rate values:

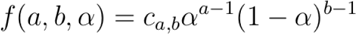

where a and b are two non-negative shape parameters. These were set to come from the uniform hyperprior distribution *a* ∼ Uniform(1, 10) and *b* ∼ Uniform(1, 10). Note that both shape parameters were assumed greater than 1, since for *a* < 1 and *b* < 1 beta distribution takes a behaviorally implausible U-shape. Precision parameter β was independently included for each submodel to provide additional flexibility in model parameter space and avoid potential dependence between competing models. Generative distribution for precision parameter *β* was represented as a lognormal distribution. Hyperpriors for lognormal parameters distributions were set to *µ* ∼ Uniform(−2.3, 3.4) for lognormal group mean and *σ* ∼ Uniform(0.01, 1.6) for lognormal standard deviation. These weakly informative hyperpriors were used in previous studies and derived from the constraints to confine group mean for the precision parameter within 0.1 to ∼ 30 interval (Meder *et al*., 2019; Nilsson *et al*., 2011). The latent-mixture part of the model assumed a subject-specific model indicator variable to represent a latent mixture of four competing submodels. This assumption allowed to model each subject response pattern independently by using a different mixture of competing submodels. Model indicator variable, z *_i_*, was modeled with uniformly distributed flat prior representing lack of expectations toward which behavioral model best explains subjects’ response patterns.

### 1.2. Markov Chain Monte Carlo

Typically, the posterior distribution *p*(*θ* | *X*) can only be calculated analytically for a limited set of simple Bayesian models. For more complex models, the analytical approach becomes ineffective and different solutions are needed. An efficient, computer-driven sampling method known as Markov Chain Monte Carlo (MCMC) has been developed to overcome this issue (Gamerman & Lopes, 2006; Kass *et al.,* 1998). The MCMC method samples posterior probability distribution *p*(*θ* | *X*) by generating Markov chains. A sufficiently large number of samples in a chain allows accurate reproduction of *p*(θ | *X*). Here, the MCMC sampling was performed using JAGS software (v4.3.0; http://mcmc-jags.sourceforge.net/) called from MATLAB R2017a via matjags.m script (v1.3.3; https://github.com/msteyvers/matjags). A sampling procedure with 2000 burn-in samples, four chains, and 15,000 samples per chain was used to estimate the hierarchical latent-mixture model.

To ensure that chain samples are unaffected by starting values and come from a stationary distribution, it is common to measure sampling convergence. A popular diagnostic measure – potential scale reduction, R^, was calculated for each model variable, and corresponding chain traceplots were visually inspected to ensure that the procedure converged. R^ combines information on the variation within and between chains to determine whether all chains reflect the same stationary target distribution. Convergence was declared for R^ values less than 1.1. For a model selection, a posterior model probability was calculated for each subject using posterior samples of the model indicator variable. PI and PD models were independently compared by marginalizing model indicator variables over subjects and calculating Bayes factors for PICI+PICD (PI model family) models versus PDCI+PDCD (PD model family) models. Bayes factor is a measure of relative evidence of two competing hypotheses calculated as:

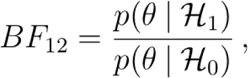

where H 1 and H 2 are two competing hypotheses. Here, these hypotheses refer to PI and PD model families generating observed behavioral data, and θ represents the model indicator variable. An analogous procedure was repeated to compare CI and CD models. A standard interpretation of Bayes factors was used to report levels of evidence (van Doorn *et al*., 2021).

The single best submodel was selected by calculating estimated model frequencies and protected exceedance probabilities (Rigoux *et al*., 2014) for the model frequency being highest among other submodels. These calculations were performed with the Variational Bayesian Analysis toolbox (v1.9.2; https://github.com/MBB-team/VBA-toolbox), which implements Bayesian model selection for group studies. Model parameters for the winning model were fitted by estimating a reduced hierarchical latent-mixture model. A full model was reduced to the single hierarchical Bayesian model by removing the model indicator variable and all branches representing competing submodels. Posterior samples for learning rates and precision were used to derive point estimates for these parameters.

### 1.3. Parameter recovery

The winning PDCI submodel was independently evaluated as a generative model for subjects’ behavioral responses. A hierarchical model comprising the branch of the full HLM model without the model indicator variable node was created and sampled using the MCMC sampling technique. Posterior distributions of learning rates and precision nodes enabled estimating individual values of these latent cognitive variables. First, group-level distributions of behavioral parameters were investigated by marginalizing individual distributions over subjects. Learning rates for positive prediction error were higher than learning rates for negative prediction error (α + =0.737 ± 0.275; α − = 0.415 ± 0.149). Bayesian hypothesis testing was used to determine whether learning rates for positive prediction errors are higher than for negative prediction errors. Moderate evidence was found in favor of this hypothesis (BF = 4.78). However, after restricting this hypothesis to the 19 subjects with most of the probability mass located over the PDCI model, very strong supporting evidence (BF = 37.90) was found. These results suggest a greater influence of positive than negative prediction errors on value estimates, especially in subjects whose behavior is best explained by the PDCI model.

### 1.4. Relationship between model parameters and behavioral performance

To better understand the behavioral significance of the observed gap between learning rates for positive and negative prediction errors, we investigated a possible relationship between the difference in learning rates and the mean number of reversals. A significant negative correlation between the two variables was found (r =−0.76; p < 0.0001), indicating that the higher difference between the learning rate for positive and negative prediction error was related to the lower reversal tendency. These results suggest that probability matching behavior is reflected by (1) a high learning rate for positive prediction errors because the subjects are more likely to repeat their choice after the previous correct choice and (2) a relatively low learning rate for negative prediction errors because the subjects tend to switch their choice after a previous incorrect choice.

**Fig. S1.**
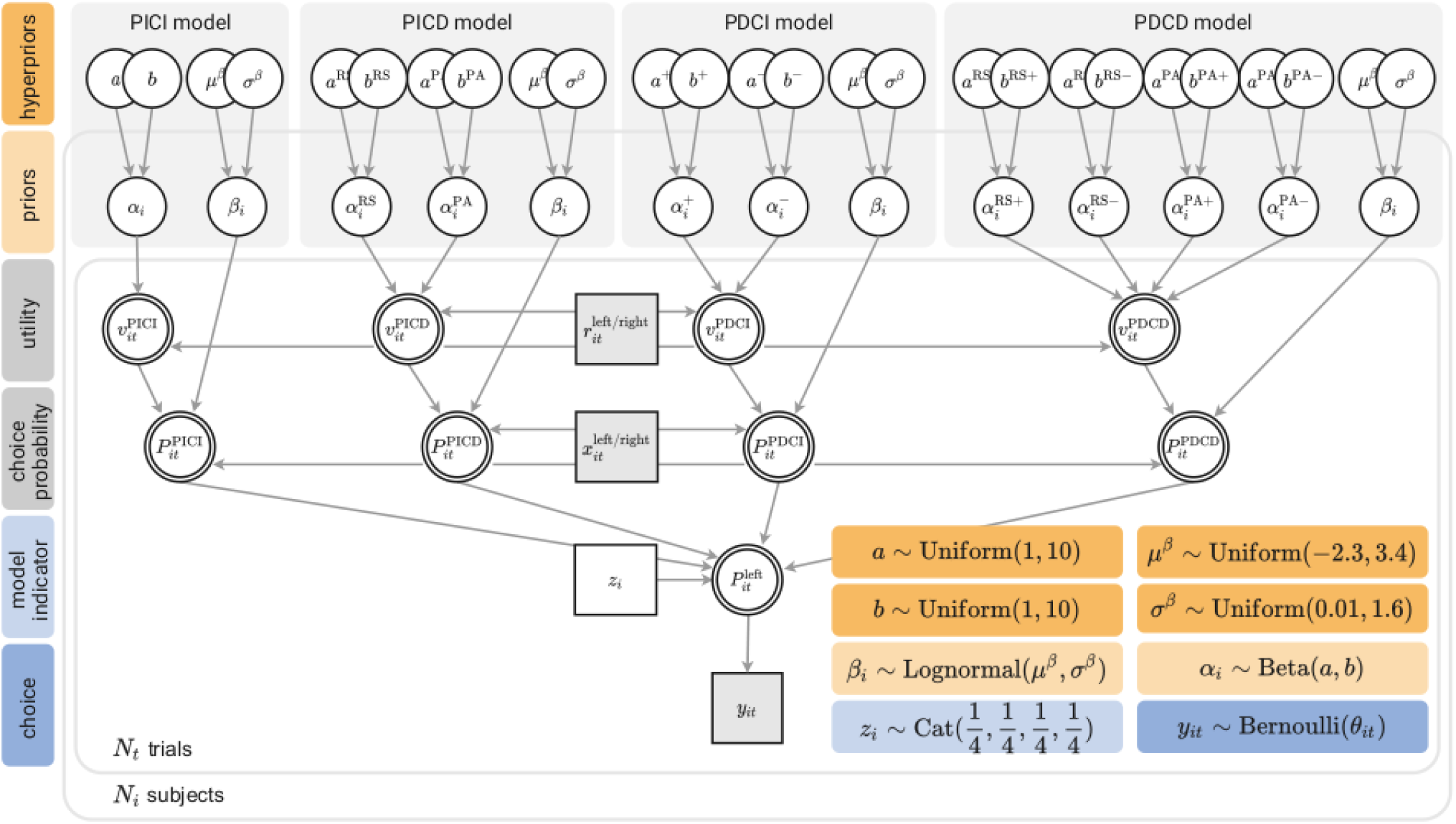
Hierarchical latent-mixture model. Choice probability for the trial *t* and subject *i* was modeled as a latent mixture of choice probabilities calculated for four different reinforcement learning submodels: PICI with single learning rate for each subject, PICD with two separate learning rates for reward-seeking and punishment-avoiding condition, PDCI with two separate learning rates for positive and negative prediction errors and PDCD with four separate learning rates for reward-seeking and punishment-avoiding conditions and positive and negative prediction errors. Probabilities for each submodel are approximated by a subject-dependent model indicator variable *z_i_*. Circular nodes represent continuous variables, square nodes represent discrete variables, unshaded nodes represent unobserved variables, shaded nodes represent observed variables, single border nodes represent stochastic variables, and double border nodes represent deterministic variables. Boxes on the left-hand side describe the role of each variable in the model, boxes on the bottom-right present the details of the prior and hyperprior distributions.

**Fig. S2.**
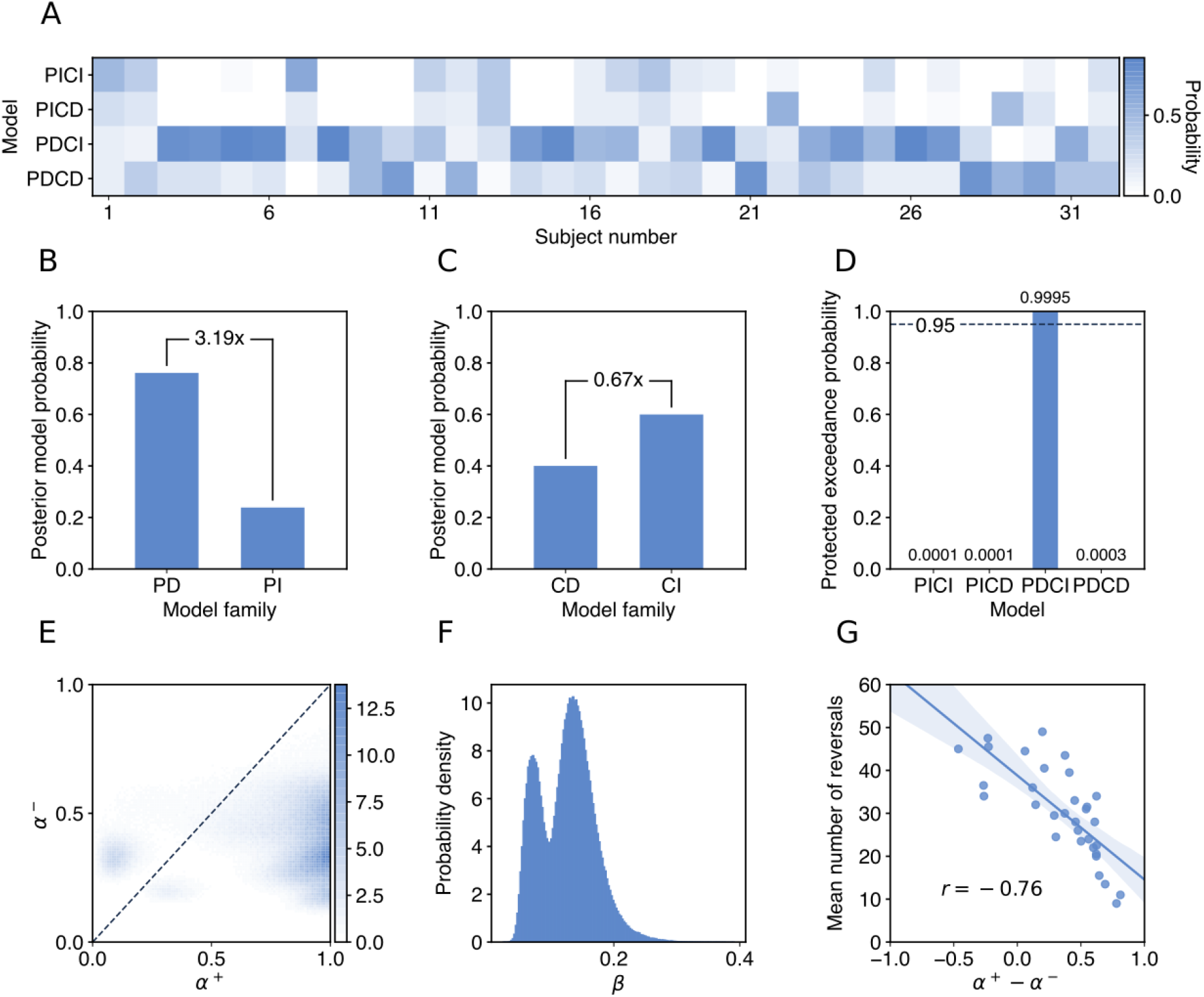
Model selection and parameter recovery results. (**A**) Posterior distribution of the model indicator variable *z_i_*. Most of the subjects had probability mass over either PDCI (19/32) or PDCD (8/32) models. (**B** and **C**) Posterior distribution of the model indicator variable *z*_i_ marginalized over subjects and model classes. Model families: prediction error dependent (PD; PDCI and PDCD models); prediction error independent (PI; PICI and PICD models); condition dependent (CD; PICD and PDCD models); condition independent (CI; PICI and PDCI models). Numbers over the bars represent Bayes factors for the hypothesis that behavioral responses are better explained by the models with separate learning rates for (**B**) positive and negative prediction errors or (**C**) reward-seeking and punishment-avoiding conditions. (**C**) Protected exceedance probabilities for each submodel being more frequent than all other submodels. (**E**) Posterior probability distribution of learning rates for positive, *α^+^*, and negative prediction errors, *α^-^*, for the winning PDCI model. The hierarchical model with PDCI submodel as a generative model for subject responses was evaluated independently of the main HLM model. (**F**) Posterior probability distribution of the precision parameter for the winning PDCI model. (**G**) Relationship between the difference in positive and negative learning rates estimated from the PDCI model and a mean number of reversals indicating probability matching behavior.

## 2. fMRI data processing

### 2.1. Preprocessing

Neuroimaging data were preprocessed using fMRIPrep version 1.4.1 (Esteban *et al*., 2019; Gorgolewski *et al*., 2011). The T1-weighted images were corrected for intensity non-uniformity (INU) with N4BiasFieldCorrection (Tustison *et al*., 2010), distributed with ANTs 2.2.0 (Avants *et al*., 2008), and used further used as reference T1-weighed image. The T1w reference was then skull-stripped with a Nipype implementation of the antsBrainExtraction.sh workflow (from ANTs), using OASIS30ANTs as the target template. Brain tissue segmentation of cerebrospinal fluid (CSF), white matter (WM), and gray matter was performed on the brain-extracted T1w using FAST (FSL 5.0.9, Zhang *et al*. (2000)). Brain surfaces were reconstructed using recon-all (FreeSurfer 6.0.1, Dale *et al*. (1999)), and the brain mask estimated previously was refined with a custom variation of the method to reconcile ANTs-derived and FreeSurfer-derived segmentation of the cortical gray matter of Mindboggle (Klein *et al*., 2017). Volume-based spatial normalization to the standard space (MNI152NLin2009cAsym) was performed through nonlinear registration with antsRegistration (ANTs 2.2.0), using brain-extracted versions of both T1w reference and the T1w template. The ICBM 152 Nonlinear Asymmetrical template was selected for spatial normalization (version 2009c; TemplateFlow ID: MNI152NLin2009cAsym; Yoon *et al*. (2009)).

For each of the two fMRI runs per subject, the following preprocessing was performed. First, a reference volume and its skull-stripped version were generated using a custom methodology of fMRIPrep. The blood-oxygen-level dependent reference was then co-registered to the T1w reference using bbregister (FreeSurfer), which implements boundary-based registration (Greve & Fischl, 2009). Co-registration was configured with nine degrees of freedom to account for distortions remaining in the BOLD reference. Head-motion parameters for the BOLD reference were estimated before any spatiotemporal filtering using mcflirt (FSL 5.0.9, (Jenkinson, 2003)). fMRI time series were slice-time corrected using 3dTshift from AFNI 20160207 (Cox & Hyde, 1997).

Then, time series were resampled to surfaces on the fsaverage5 space, their original native space, and standard MNI152NLin2009cAsym space by applying a single composite transform to correct head motion and susceptibility distortions. These resampled time series are referred to as preprocessed time series. First, a reference volume and its skull-stripped version were generated using a custom methodology of fMRIPrep. Several confounding time series were calculated based on the preprocessed time series: framewise displacement (FD), a spatial standard deviation of successive difference images (DVARS), and three region-wise global signals. FD and DVARS were estimated for each functional run, using their implementations in Nipype. The three global signals were extracted from the CSF, WM, and whole-brain masks. Additionally, head-motion estimates calculated during the correction step were stored as confound time series. All head-motion and global signal confounds were expanded with the inclusion of temporal derivatives and quadratic terms (Satterthwaite *et al*., 2013). Frames that exceeded a threshold of 0.5 mm FD or 1.5 standardized DVARS were marked as motion outliers.

### 2.2. Model-based fMRI

#### 2.2.1 First-level GLM

The single-subject effect of processing increasing prediction errors was modeled with a subject-level general linear model. Two separate GLMs were created for reward-seeking and punishment-avoiding task conditions. Each GLM was composed of regressors modeling task events and estimated latent decision parameters and nuisance regressors accounting for noise components related to head motion and scanner drift. Task regressors of interest included regressor modeling onset of the outcome phase and its parametric modulation with the estimated trial-wise prediction error (PE regressor). Task regressors of no interest modeled other experimental events: onset of the decision and its parametric modulation with the expected probability that chosen option will be correct, left and right button press, trials with a missing response, offset of the outcome phase (**Fig. S3**). Nuisance regressors included 24 head motion parameters to remove residual movement artifacts and cosine functions derived from the cosine drift model to remove low-frequency artifacts. All task events were modeled as impulse functions with zero duration and convolved with a canonical double-gamma hemodynamic response function. First-level contrast was defined as a one-sample t-test for the PE regressor effect against the baseline. Resulting statistical parametric maps were used in the second level of analysis.

**Fig. S3.**
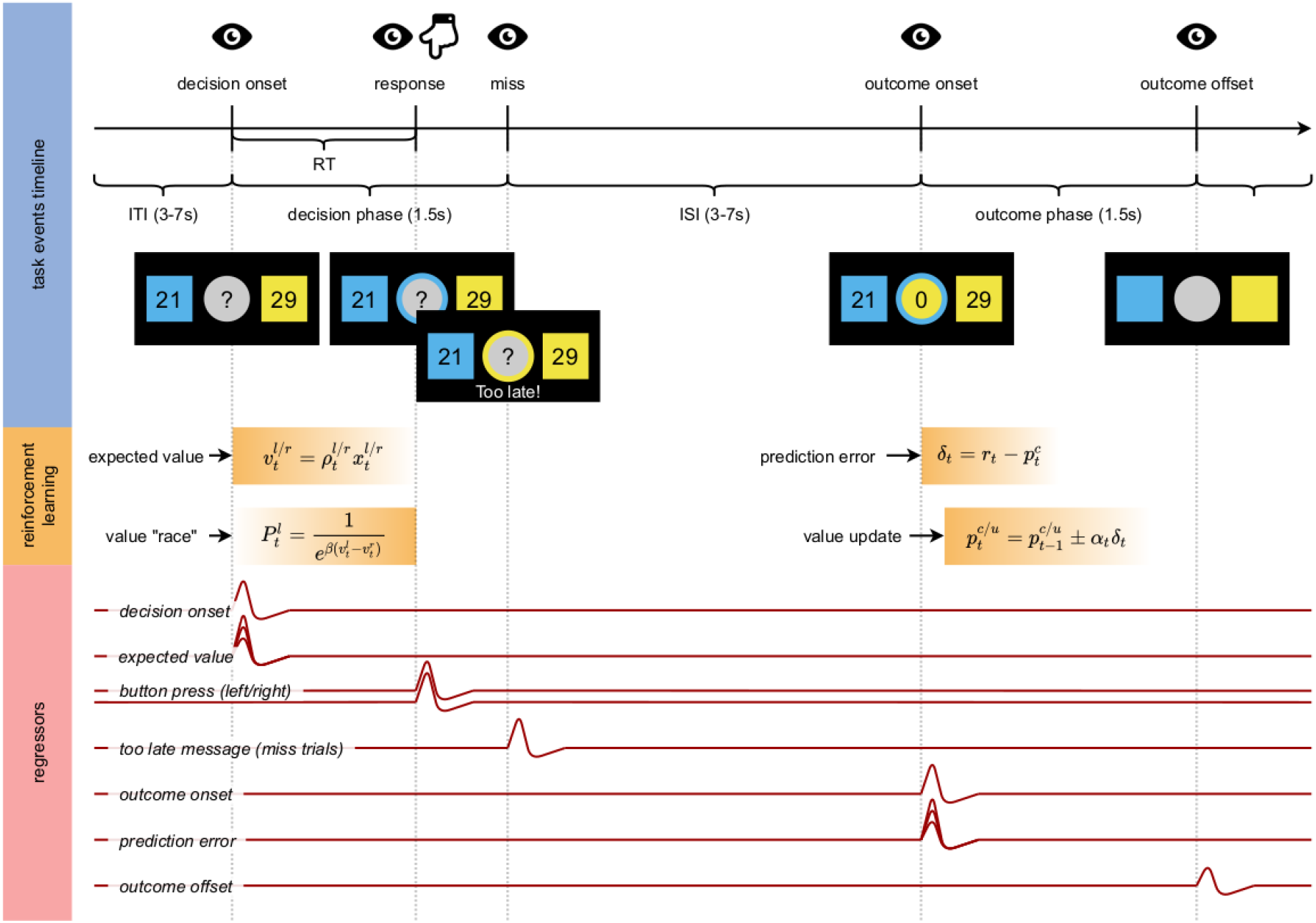
Event model for a single trial. Task events, latent computational variables, and GLM regressors are shown on a timeline for a single experimental trial. **The top panel** shows a trial timeline with corresponding visual changes (eye icon) and motor responses (hand icon). **The middle pane**l shows latent reinforcement learning variables like expected value and prediction errors time-locked to task events. **The bottom panel** presents modeled BOLD responses used in first-level GLM.

#### 2.2.1 Second-level GLM

The second level of the analysis was performed as a random-effects analysis separately modeling each task condition and within-subject effects. The significance of the two effects of interest across the group was tested: (1) the combined effect of increasing and decreasing prediction error across both task conditions and (2) the difference in response to increasing and decreasing prediction error between reward-seeking and punishment-avoiding conditions. A threshold of *p* < 0.0001 with false discovery rate (FDR) correction was used for the former effect, accounting for its high statistical power compared with the latter effect. This stringent thresholding was imposed to reduce the number of significant clusters and improve the readability of the statistical maps. A threshold of *p* < 0.001 with FDR correction was used for the latter effect. An additional threshold for a cluster size of 20 connected voxels was used in either case.

Differences in response to PE between reward-seeking and punishment-avoiding conditions cannot be unambiguously interpreted since statistical testing of this effect relies on testing the difference between slopes of the regression between voxel activity and the prediction error. For example, a significant response for increasing PEs in reward-seeking compared to the punishment-avoiding condition may be related to one of the three effects: (1) for both conditions, activity is correlated with increasing PE, but for the reward-seeking condition the relationship is significantly stronger, (2) for both conditions activity is correlated with decreasing PE, but for reward-seeking condition, the relationship is significantly weaker or (3) activity is correlated with increasing PE in reward-seeking condition and correlated with decreasing PE in punishment-avoiding condition. A post hoc test on the cluster level was performed to discriminate between these effects. Response profiles were extracted for each subject, task condition, and significant cluster exhibiting increased or decreased response to prediction error in reward-seeking condition compared to punishment-avoiding condition. These profiles were calculated as first-level parameter estimates, i.e., GLM beta values, for peak voxel for the PE regressor effect. Then, average beta values across subjects were calculated for both conditions. To further confirm the significant change in response profiles, a paired t-test between reward-seeking and punishment-avoiding betas was performed for all clusters.

#### 2.2.2 Context-independent prediction error processing

We wanted to identify brain regions in which activity increases with increasing PEs and decreases with increasing PEs. We used individually calculated prediction errors to construct PE regressors modeling latent PE computation time-locked to the outcome onset. Our first aim was to identify brain regions encoding increasing and decreasing PEs regardless of the task condition. These regions form the core of the learning network enabling the processing of positive and negative feedback information used to update the values of chosen stimuli. A broad network of regions in which activity was correlated with increasing and decreasing PEs was found (**Fig. S4**). For increasing PEs, significant activity was found in clusters in the vmPFC, superior temporal gyrus, bilateral orbitofrontal cortex, bilateral putamen, left postcentral gyrus, right amygdala, and VS, left posterior cingulate cortex (PCC), left angular gyrus, right precentral gyrus, left middle temporal gyrus (**Table S1**; increasing PE). Other significant clusters were located in the left caudal middle frontal gyrus, bilateral planum temporale, left parahippocampal gyrus, and right superior frontal gyrus. For decreasing PEs, significant activity was found in clusters in the left dorsomedial cingulate cortex, bilateral anterior insula (aINS), right pars opercularis, left dorsolateral prefrontal cortex (dlPFC), left cerebellum, left precentral gyrus, and left superior frontal gyrus (**Table S1**; decreasing PE). Generally, activity related to increasing PEs was more widespread than activity related to decreasing PEs.

**Fig. S4.**
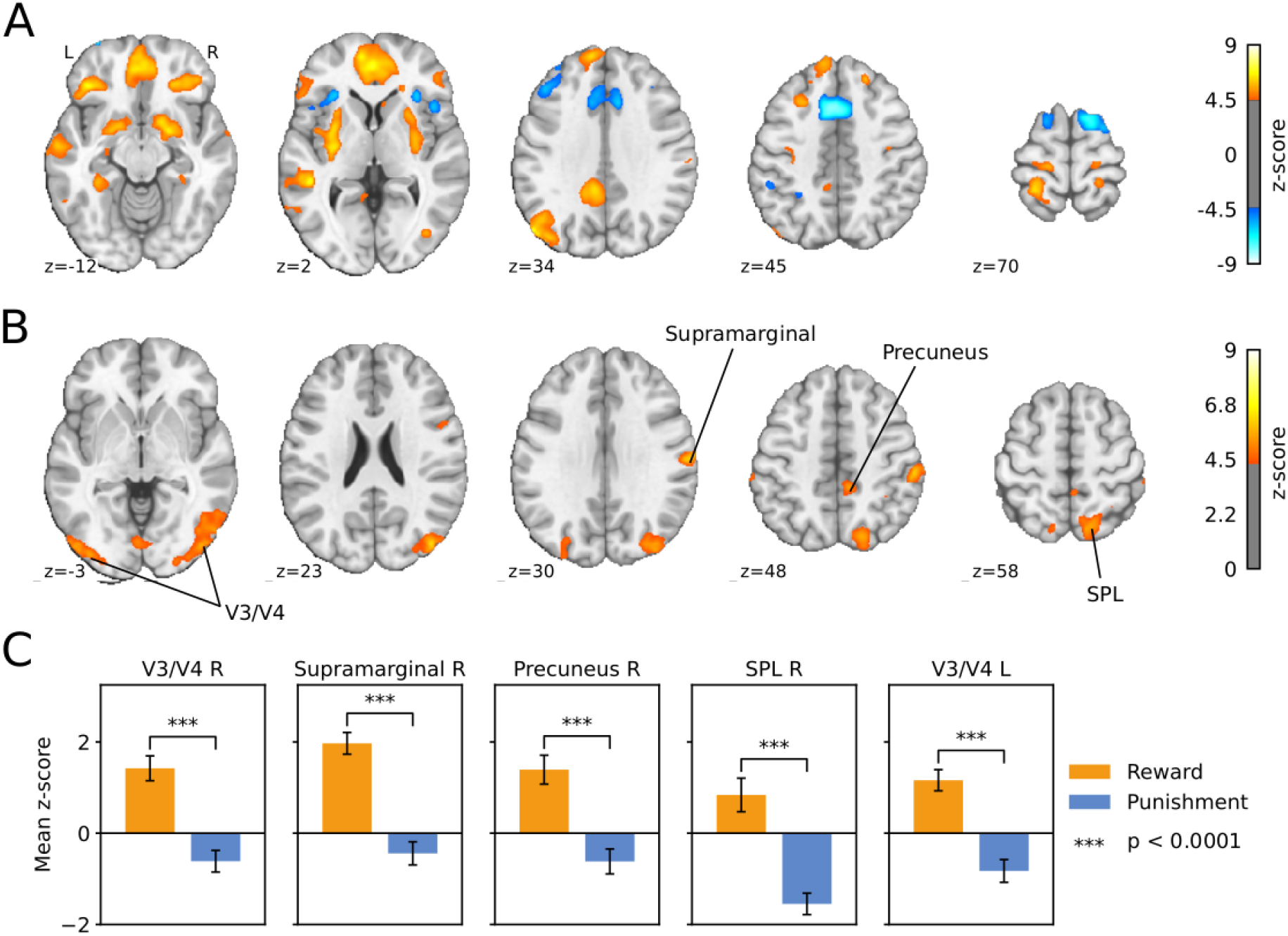
Model-based fMRI results. (**A**) Brain regions processing increasing and decreasing prediction errors. Orange regions exhibit increased activity with increasing prediction error in both task conditions. Blue regions decrease their BOLD response with increasing prediction error in both task conditions. Statistical maps were FDR corrected with a threshold p < 0.0001. (**B**) Brain regions responding differently to prediction errors in rewards-seeking and punishment-avoiding conditions. For orange clusters slope between voxel activity and PE was higher in reward-seeking compared to punishment-avoiding conditions. (**C**) Post-hoc test for the clusters in B (only five clusters with the highest z-scores are shown). First-level parameter estimates (z-scores) for the PE regressor were extracted individually for each cluster, subject, and condition. Each region in B increased its activity with increasing PE in the reward-seeking condition and decreased its activity with increasing PE in the punishment-avoiding condition (paired t-test; p<0.0001). Abbreviations: SPL - superior temporal lobule.

#### 2.2.3 Context-dependent prediction error processing

We wanted to test whether PE signaling in essential parts of an increasing PE network like striatum and vmPFC undergo valence-related changes. To investigate differences in PE processing between task conditions, we evaluated a second-level GLM testing for higher response to increasing PEs in reward-seeking than punishment-avoiding conditions. A set of brain regions was found for which the regression slope between voxel activity and the prediction error was higher in the reward-seeking compared to punishment-avoiding condition (**Fig. S4B**). These regions included bilateral visual areas V3 and V4, right supramarginal gyrus, right superior parietal lobule, right precuneus, right primary visual cortex, and right precentral gyrus (Table 4.2). None of these regions were part of the core PE processing network. Since differences in response to PE between reward-seeking and punishment-avoiding conditions cannot be unequivocally interpreted, a posthoc test was performed for all significant clusters by extracting individual subject and condition response profiles as first-level parameter estimates for the peak voxel. Positive activity scaling with increasing PEs in reward-seeking condition and negative activity scaling with increasing PEs in punishment-avoidance condition was found in all seven clusters (paired t-test; *p*<0.0001; **Fig. S4C**). There were no brain regions with significantly higher responses to increasing PE in punishment-avoiding condition.

**Table S1.** Between-condition differences in PE signaling. Regions exhibiting differences in prediction error processing between reward-seeking and punishment-avoiding conditions.

| Region | L/R | X | Y | Z | Z-score peak | Cluster Size (mm<sup>3</sup>) |
| --- | --- | --- | --- | --- | --- | --- |
| Increasing PE |  |  |  |  |  |  |
| Ventromedial Prefrontal Cortex | L | -3 | 48 | -1 | 7.23 | 30397 |
| Superior Temporal Gyrus | L | -48 | -39 | 3 | 6.93 | 1417 |
| Lateral Orbitofrontal Cortex | L | -39 | 33 | -12 | 6.79 | 5071 |
| Putamen | L | -30 | -12 | 6 | 6.63 | 8473 |
| Postcentral Gyrus | L | -27 | -42 | 66 | 6.62 | 1732 |
| Amygdala, Ventral Striatum, Putamen | R | 21 | 0 | -12 | 6.52 | 8347 |
| Posterior Cingulate Cortex | L | -6 | -45 | 38 | 6.47 | 9985 |
| Angular Gyrus | L | -54 | -75 | 34 | 6.32 | 9009 |
| Precentral Gyrus | R | 24 | -24 | 59 | 6.30 | 2803 |
| Middle Temporal Gyrus | L | -60 | -12 | -15 | 6.18 | 2425 |
| Caudal Middle Frontal Gyrus | L | -27 | 24 | 48 | 6.07 | 1039 |
| Lateral Orbitofrontal Cortex | R | 27 | 36 | -12 | 6.00 | 3213 |
| Planum Temporale, Supramarginal Gyrus | R | 45 | -36 | 17 | 5.86 | 3244 |
| Parahippocampal Gyrus | L | -33 | -39 | -12 | 5.80 | 1417 |
| Precentral Gyrus | L | -15 | -27 | 73 | 5.69 | 1039 |
| Superior Frontal Gyrus | R | 21 | 39 | 55 | 5.69 | 1386 |
| Planum Temporale | L | -57 | -33 | 17 | 5.69 | 1669 |
| Middle Temporal Gyrus | L | -57 | -57 | -1 | 5.39 | 850 |
| Precentral Gyrus | L | -6 | -18 | 52 | 5.39 | 787 |
| Postcentral Gyrus | L | -39 | -24 | 52 | 5.28 | 913 |
| Decreasing PE |  |  |  |  |  |  |
| Dorsomedial Cingulate Cortex, SMA | L | -6 | 12 | 48 | 8.18 | 14647 |
| Anterior Insula | L | -33 | 21 | 10 | 6.71 | 1543 |
| Anterior Insula | R | 30 | 24 | 6 | 6.43 | 756 |
| Pars Opercularis | R | 45 | 18 | 3 | 5.93 | 787 |
| Dorsolateral Prefrontal Cortex | L | -45 | 30 | 34 | 5.89 | 2803 |
| Cerebellum | L | -33 | -60 | -26 | 5.81 | 630 |
| Precentral Gyrus | L | -30 | -3 | 59 | 5.62 | 913 |
| Superior Frontal Gyrus | L | -18 | 6 | 69 | 5.55 | 882 |

**Table S2.**
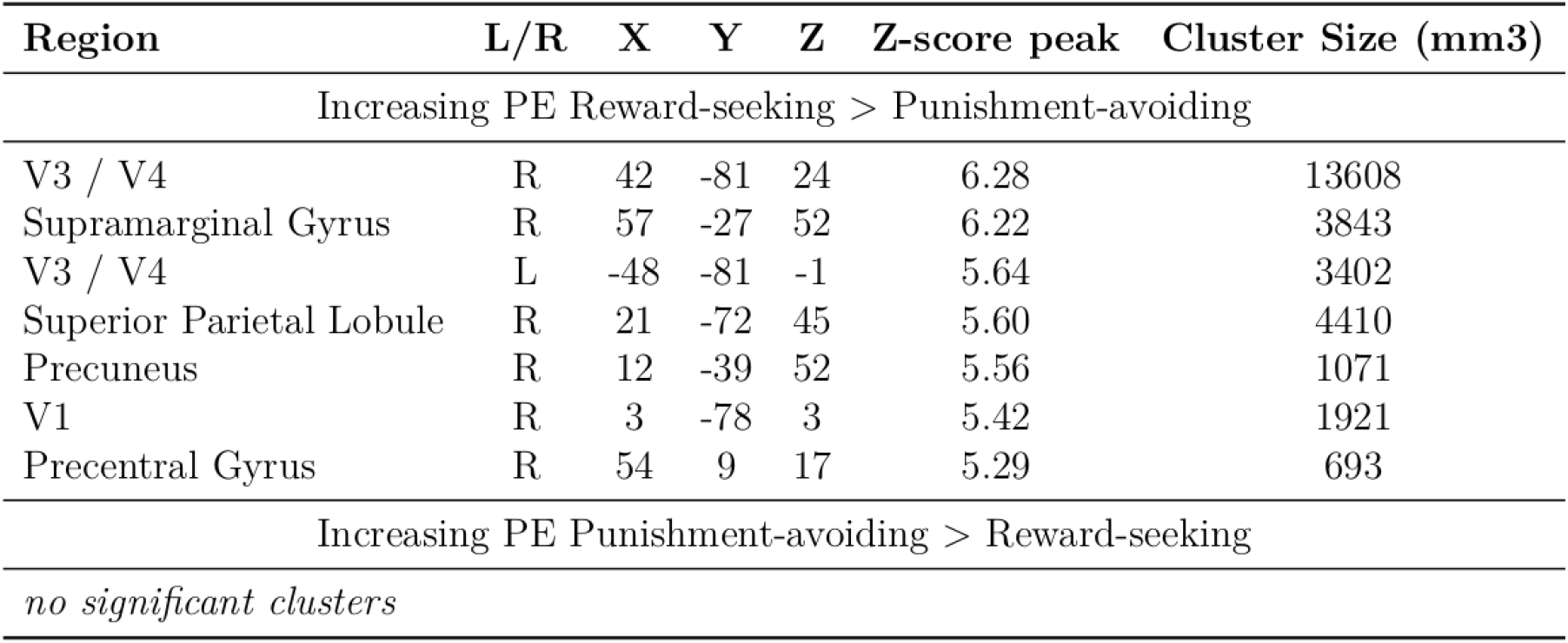
Between-condition differences in PE signaling. Regions exhibiting differences in prediction error processing between reward-seeking and punishment-avoiding conditions.

| Region | L/R | X | Y | Z | Z-score peak | Cluster Size (mm3) |
| --- | --- | --- | --- | --- | --- | --- |
| Increasing PE Reward-seeking > Punishment-avoiding |  |  |  |  |  |  |
| V3 / V4 | R | 42 | -81 | 24 | 6.28 | 13608 |
| Supramarginal Gyrus | R | 57 | -27 | 52 | 6.22 | 3843 |
| V3 / V4 | L | -48 | -81 | -1 | 5.64 | 3402 |
| Superior Parietal Lobule | R | 21 | -72 | 45 | 5.60 | 4410 |
| Precuneus | R | 12 | -39 | 52 | 5.56 | 1071 |
| V1 | R | 3 | -78 | 3 | 5.42 | 1921 |
| Precentral Gyrus | R | 54 | 9 | 17 | 5.29 | 693 |
| Increasing PE Punishment-avoiding > Reward-seeking |  |  |  |  |  |  |
| *no significant clusters* |  |  |  |  |  |  |

### 2.3. Analysis of functional brain networks

#### 2.3.1 Brain parcellation

One of the first steps in brain network analysis is dividing the brain into regions that will be considered nodes of the functional network (Sporns, 2013). Each specific division is called brain parcellation or brain atlas. The frequently used brain parcellation in resting-state and task-based fMRI studies is the parcellation introduced by Power *et al*. (2011). It consists of 264 regions of interest defined based on a meta-analysis of activation patterns from multiple studies. Each ROI is modeled as a 5mm sphere around the specific MNI coordinates. Power’s parcellation additionally includes the division of the 264 ROIs into 13 large-scale networks: auditory, cerebellar, somatomotor, cingulo-opercular, default mode, memory, ventral attention, dorsal attention, fronto-parietal, salience, subcortical, uncertain and visual. This division allows examining the dynamic interplay between LSNs in various experimental conditions. However, Power’s parcellation does not contain separate prediction-error-related networks; therefore, it cannot be directly used to study how the reward system interacts with the rest of the brain during reinforcement learning. To overcome this issue, the Power’s ROIs were combined with regions from the recent meta-analysis on neural representations of prediction error valence (Fouragnan *et al*., 2018). This procedure allows studying effects specific for networks signaling prediction errors while still preserving reference to well-known large-scale brain systems. Moreover, a recent meta-analysis provided evidence that reward-related brain regions form a stable network observed even during rest (Huckins *et al*., 2019).

The following procedure was employed to create extended brain parcellation. First, two prediction-error-related networks were created based on cluster centers for the valence analysis in a recent meta-analysis on neural correlates of prediction errors (Table 2 in Fouragnan *et al*. (2018)). The network signaling positive prediction error (+PE) was created from regions exhibiting higher BOLD activation for positive than negative prediction errors (pattern A (ii) in Fouragnan *et al*. (2018)). Conversely, the network signaling negative prediction error (−PE) consisted of regions with a higher BOLD response for negative than positive prediction errors (pattern A(i) in Fouragnan *et al*. (2018)). Second, for each cluster center, a new ROI was created using one of the three strategies based on its distance to the closest ROI in Power’s atlas (d):

- If d > 10mm, there was no overlap between the 5mm sphere created at the cluster center and existing ROIs, so a new ROI was created and added to the ROIs set without any modification of the existing base parcellation.
- If 10mm > d > 5mm, there was a slight overlap between the 5mm sphere created at the cluster center and the closest Power’s ROI (no more than 31% of sphere volume overlapped). In this case, the closest Power’s ROI was shifted into the location of the ALE cluster center and renamed according to the assignment to one of the prediction-error-related networks.
- If d < 5mm there was a significant overlap between the 5mm sphere created at the cluster center and the closest Power’s ROI (more than 31% of sphere volume overlapped). In this case, the original Power’s ROI was retained at its original location and relabelled according to the assignment to one of the prediction-error-related networks.

All newly created ROIs were initially modeled as 5mm spheres around specific MNI coordinates. Radi of four striatal ROIs – left and right pallidum and left and right ventral striatum were decreased to 4mm to avoid overlap between spheres (**Table S3**). The extended brain parcellation consisted of 272 ROIs divided into 15 large-scale networks (13 LSNs from Power *et al*. (2011) and 2 PE networks). Four ROIs from the uncertain network – MNI coordinates (-31, -10, -36), (-56, -45, -24), (8, 41, -24), and (52, -34, -27) – were further excluded due to signal dropout in some participants. The final parcellation consisted of N ROI = 268 regions of interest.

**Table S3.** Between-condition differences in PE signaling. Prediction error signaling ROIs created from ALE clusters from Fouragnan et al. (2018).

| Region | R/L | X | Y | Z | r(mm) | Original network | Strategy |
| --- | --- | --- | --- | --- | --- | --- | --- |
| ROIs signaling positive prediction errors (+PE) |  |  |  |  |  |  |  |
| Ventrolateral orbitofrontal cortex | R | 32 | 44 | -10 | 5 | Uncertain | shifted |
| Ventral striatum | L | -12 | 8 | -4 | 4 | - | created |
| Posterior cingulate cortex | L | 0 | -36 | 26 | 5 | Memory | shifted |
| Medial prefrontal cortex | L | -2 | 46 | 20 | 5 | Default mode | shifted |
| Ventral striatum | R | 8 | 8 | -2 | 4 | - | created |
| Dorsomedial prefrontal cortex | L | -6 | -56 | 14 | 5 | Default mode | shifted |
| Ventromedial prefrontal cortex | L | -2 | 42 | 0 | 5 | Default mode | shifted |
| ROIs signaling negative prediction errors (-PE) |  |  |  |  |  |  |  |
| Anterior insula | L | -32 | 22 | -4 | 5 | Salience | shifted |
| Middle frontal gyrus | R | 32 | 14 | 56 | 5 | Fronto-parietal | relabelled |
| Inferior parietal lobule | R | 40 | -48 | 42 | 5 | Fronto-parietal | shifted |
| Superior temporal sulcus | R | 54 | -43 | 22 | 5 | Ventral attention | relabelled |
| Fusiform area | L | -42 | -60 | -9 | 5 | Dorsal attention | relabelled |
| Pallidum | R | 12 | 8 | 4 | 4 | Subcortical | shifted |
| Dorsolateral prefrontal cortex | R | 40 | 34 | 30 | 5 | Salience | shifted |
| Dorsolateral prefrontal cortex | L | -42 | 25 | 30 | 5 | Fronto-parietal | relabelled |
| Middle temporal gyrus | R | 60 | -28 | -6 | 5 | Default mode | shifted |
| Inferior parietal lobule | L | -38 | -48 | 42 | 5 | Dorsal attention | shifted |
| Thalamus | L | -6 | -26 | 8 | 5 | Subcortical | shifted |
| Anterior insula | R | 32 | 24 | -2 | 5 | Salience | shifted |
| Pallidum | L | -14 | 6 | 2 | 4 | Subcortical | shifted |
| Dorsomedial prefrontal cortex | R | 20 | 50 | 4 | 5 | - | created |
| Dorsomedial orbitofrontal cortex | R | 38 | 58 | -2 | 5 | - | created |
| Precentral cortex | L | -52 | 0 | 34 | 5 | - | created |
| Habenula | R | 2 | -20 | -18 | 5 | - | created |
| Amygdala | R | 18 | -6 | -12 | 5 | - | created |
| Middle frontal gyrus | R | 38 | 4 | 32 | 5 | - | created |
| Middle frontal gyrus | L | -28 | 12 | 60 | 5 | Fronto-parietal | shifted |
| Dorsomedial cingulate cortex | R | 5 | 23 | 37 | 5 | Salience | relabelled |

#### 2.3.2 Beta-series correlation

Task-related functional connectivity was estimated using a beta-series correlation method introduced by Rissman *et al*. (2004). The method’s name comes from the GLM parameter beta representing the linear coefficient reflecting the contribution of modeled signal into an observed BOLD response. The BSC method quantifies event-to-event fluctuations in the activity of different brain areas. These fluctuations allow for estimating statistical dependency between regional activations for different event types.

In BSC, each trial is modeled separately in GLM. Then, a series of beta maps representing brain activations for a series of events was used to calculate condition-specific connectivity evoked by the task. An alternative methodology to calculate task-evoked functional connectivity is psychophysiological interaction analysis (Friston *et al*., 1997). However, it has been suggested that the BSC method can have more statistical power than the PPI method when applied to event-related designs with many trials and short event durations (Cisler *et al*., 2014). The “single-trial-versus-other-trial” approach introduced by Mumford *et al*. (2012) was used to compute activation beta maps. This approach assumes creating a separate GLM for each trial in which the trial of interest is modeled as a separate regressor, and all other trials are modeled as a single nuisance regressor. Simulations have shown that the “single-trial-versus-other-trial” approach produces more accurate estimates of trial-wise activation patterns (Mumford *et al*., 2012).

For each trial, a separate GLM was created. The GLM consisted of:

- One regressor modeling a trial of interest as an event of duration 0s (typically used to model very short cognitive processes) time-locked to the onset of the outcome phase, convolved with a standard SPM hemodynamic response function (HRF).
- Two regressors modeling the rest of the trials of interest as events of duration 0s time-locked to the onset of the outcome phase convolved with an HRF. One regressor modeled trials with positive prediction errors (gain in reward-seeking condition or no-loss in punishment-avoiding condition). The other modeled trials with negative prediction errors (no-gain in the reward-seeking condition or loss in the punishment-avoiding condition).
- Six regressors modeling the events of no interest: decision onset, decision onset modulated with expected probability for side for being correct, decision onset for missed trials, left button press, right button press, and decision offset for missed trials.
- Twenty-four head motion parameter regressors modeling motion-related noise
- Cosine functions modeling low-frequency scanner drift.

The GLM was high-pass filtered with the cut-off frequency of 128 Hz and fitted to participants’ data using `FirstLevelModel` class from the Nistats package. The output of this procedure consisted of a three-dimensional beta map for each trial, indicating the voxel-wise estimate of neuronal activity evoked during this trial (**Fig. S.5**). After GLM estimation, beta values for each trial were averaged in each of 268 ROIs producing 268 beta-series per subject and task condition. Beta-series were then z-transformed and separated by the sign of the prediction error of each trial. This procedure gave four beta-series for each participant: +PE trials in the reward-seeking condition, −PE trials in the reward-seeking condition, +PE trials in the punishment-avoiding condition, and −PE trials in the punishment-avoiding condition. Beta-series were then correlated using Pearson’s correlation coefficient, generating an *N ROI ×N ROI* symmetrical correlation matrix for each subject, task condition, and trial-wise prediction error sign. Correlation values were then converted to z-scores using Fisher’s z-transform. Transformed matrices represented the final subject-level estimate of the functional connectivity pattern evoked by the specific task condition and type of trial.

**Fig. S5.**
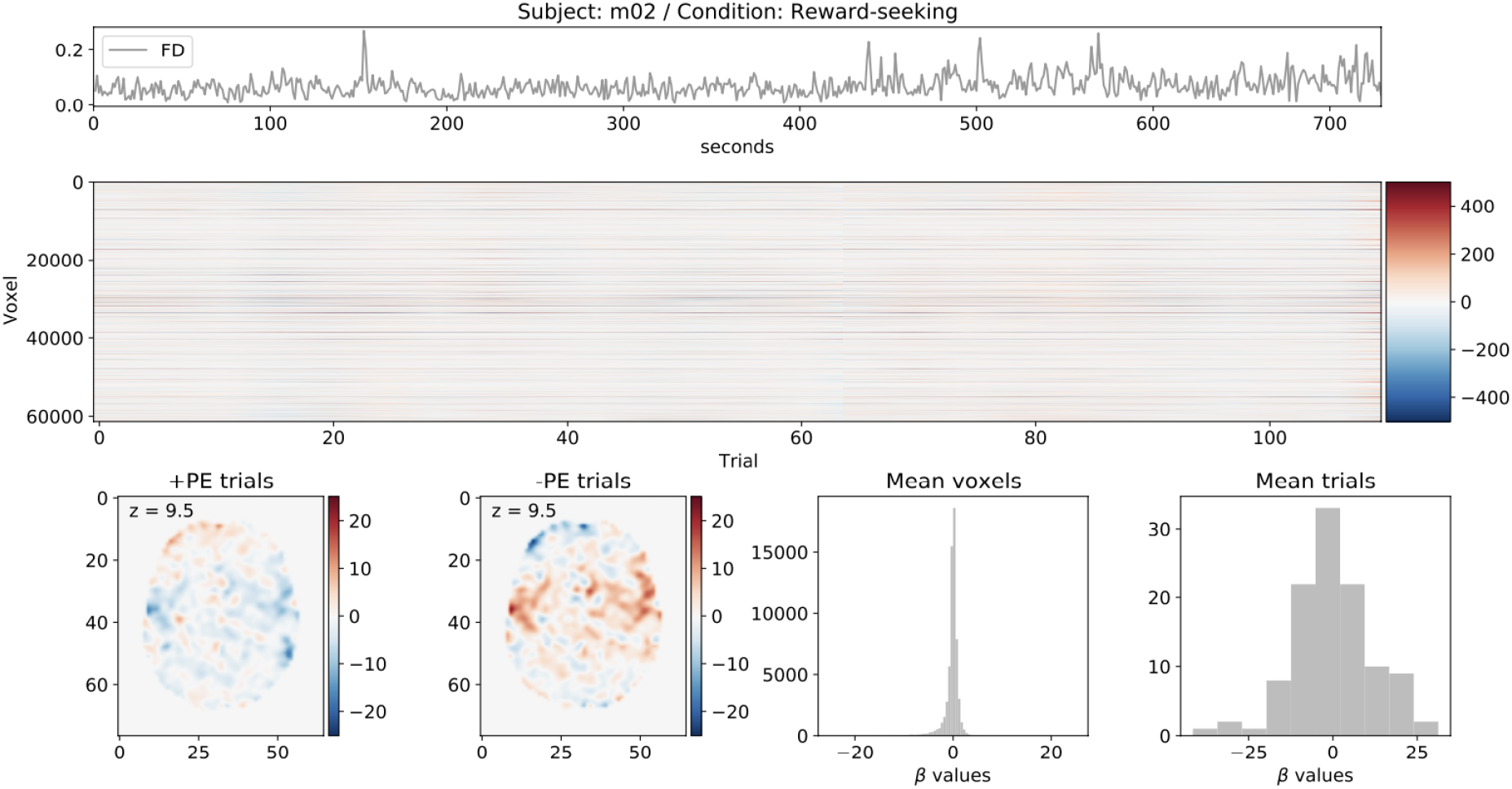
Example beta maps summary for single subject and task condition. Different characteristics of trial-wise beta maps in BSC analysis. The top panel shows the timecourse of framewise displacement FD. The middle panel shows the carpet plot of beta estimates – x-axis corresponds to 110 trials of the PRL task, y-axis corresponds to all voxels within the brain mask, color indicates trial-evoked activation. Two bottom-left panels show a coronal slice for z=9.5 for mean brain activation during +PE and -PE trials. Two bottom-right panels show beta-value distributions averaged across trials and voxels.

#### 2.3.3 Multi-scale modularity

Multi-scale modularity analysis was performed by introducing a structural resolution parameter, γ, into the modularity function:

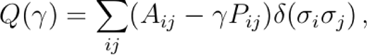

where A_ij_ is the connection strength between nodes *i* and *j*, P_ij_ is the expected connection strength according to the appropriate null model, σ_i_ indexes the community to which node *i* is assigned, and *δ* is the Kronecker delta function. Tuning the structural resolution parameter allows uncovering communities of different sizes, otherwise mathematically undetectable using standard modularity function (Fortunato & Barthelemy, 2007; Reichardt & Bornholdt, 2006). Specifically, lower values of γ result in fewer larger communities, and higher values produce more and smaller communities. At the lower end of the γ range, all network nodes are placed within one community, whereas at the higher end, there are as many communities as there are nodes. Different heuristics have been proposed to find an appropriate sampling of structural resolution parameter space. For example, similarity measures between partitions have been used to minimize the variability in the results of modularity maximization techniques (Doron *et al*., 2012; Lancichinetti *et al*., 2009). This choice has been guided by the intention to describe network reorganization with reference to well-known LSNs. Two constraints have been imposed on the analyzed topological scales:

1. Number of detected modules and mean module size should be similar to the reference partition. In other words, large γ values generating partitions with many singletons, i.e., single node communities should be excluded from the analysis.
2. Detected communities should exhibit high similarity to the reference partition while preserving a high degree of inter-individual variability. High similarity to the reference partition justifies the interpretation of the results in terms of interactions between a priori LSNs. High inter-individual variability ensures that the analysis will be more sensitive to topological changes between task conditions and prediction error signs.

To calculate community structure properties across topological scales, the structural resolution parameter space was sampled by varying γ from 0.05 to 3 with a step of 0.05. For each value of γ, the Louvain modularity maximization algorithm was used to quantify the optimal network structure (Blondel *et al*., 2008). Negative connections were separately included in the modularity function and treated asymmetrically as suggested in Rubinov and Sporns (2011):

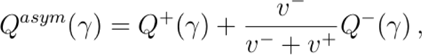

where Q ± (γ) are modularity values calculated separately for positive and negative connections, and v ± = ij A ± ij are total connection strengths, i.e., costs for positive and negative connections. For each functional network, the Louvain algorithm was run 100 times, and the partition with the highest value of *Q asym* (γ) was kept for further analysis. Several metrics were computed to inspect different topological scales (**Fig. S6**): (1) a mean number of communities, singletons, and non-singletons (communities with more than one node), (2) mean community size, (3) mean similarity between data-driven partitions, and (4) mean similarity between reference partition and data-driven partitions. Partition similarity was computed as the normalized mutual information between community partition vectors (Meilă, 2007).

The mean number of data-driven communities was closest to 14 reference communities for γ = 1.5 (average 14.59 communities per partition). For γ > 1.5, there was a steep increase in the number of singletons, whereas the number of non-singletons rose to 21.85 for γ = 2 and plateaued with a further increase in structural resolution. The average data-driven community size was above 100 nodes per partition for 0 < γ < 0.9 and rapidly declined for γ ∼ 1. For γ = 1.65, the average community size was closest to the average reference community size (18.86 nodes per community for data-driven partitions; 19.14 nodes per community for reference partition). The similarity between partitions measured as normalized mutual information increased as a function of γ. For 0.15 ≤ γ ≤ 2.1, data-driven partitions exhibited higher similarity to themselves than to reference partition. The ratio between data-driven partition similarity, M*I_d−d_*, and similarity between reference and data-driven partition, M*I_d−d_*= 1.85). *M I_d−r_*, was highest for γ = 0.4.

According to the first heuristic, the analyzed γ range should include structural resolutions for which the number of communities and the average size of the community resembles characteristics of the reference partition. This consideration suggests including γ ∼ 1.5 since the size and number of data-driven communities were closest to the size and number of reference communities for this topological scale. On the other hand, the second heuristic suggests including γ ∼ 0.5 because partitions observed for this topological scale exhibit the optimal similarity to the reference partition while preserving a high degree of inter-individual variability. For γ > 2 number of singletons starts to dominate non-singleton communities, and relative inter-individual variability decrease compared to similarity to reference partition, suggesting that this part of the structural resolution scale should be excluded from further analysis. Considering both heuristics, the final range 0.5 ≤ γ ≤ 1.5 was adopted for all subsequent analyses. Whereas dense sampling of this range (e.g., with a step of 0.05) would allow tracing how observed effects smoothly appear and disappear when changing a topological resolution, it would also be computationally expensive and challenging to describe statistically. Therefore, a sparse sampling of γ range was employed, enabling to focus on three distinct topological scales: γ = 0.5 representing the largest scale of few “super-communities” consisting of multiple LSNs, γ = 1 representing the most frequently studied intermediate scale, and γ = 1.5 representing finer scale approximately resembling well-known division into LSNs.

**Fig. S6.**
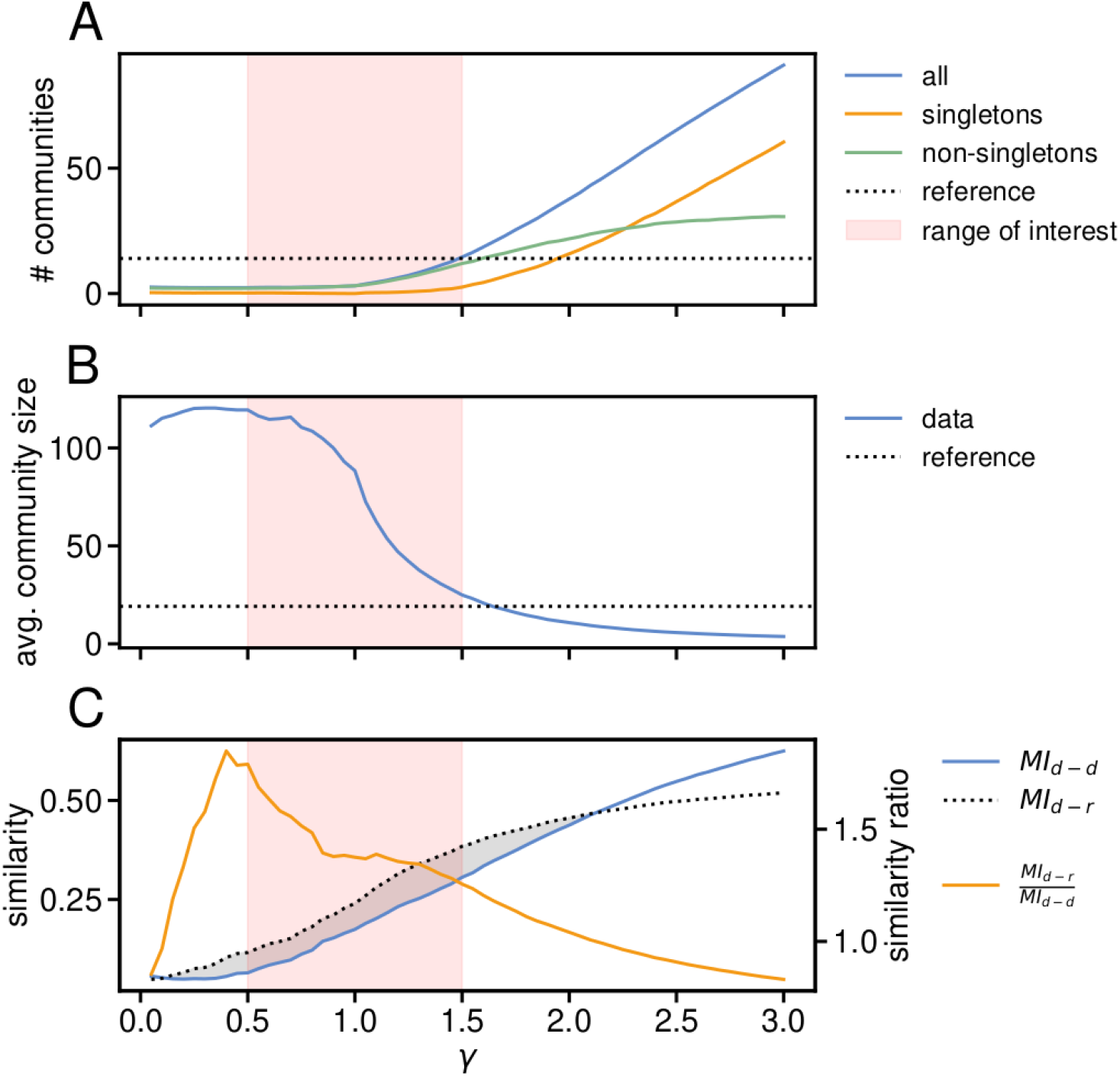
Structural resolution parameter analysis. The structural resolution parameter, γ, was varied from 0.05 to 3 with a step of 0.05. For each value of γ, optimal community structures for empirical networks were calculated using 100 repetitions of the Louvain algorithm (Blondel et al., 2008). (**A**) The number of all communities, singleton communities, and non-singleton communities. Reference partition based on Power ROIs described in section 4.6.1 has 14 communities. (**B**) Mean data-driven community size compared with the average size of 19.14 ROIs per community for reference partition. (**C**) Average similarity between data-driven partitions and between reference and data-driven partitions. The gray area indicates γ range for which partitions are more similar to the reference than to themselves. The orange line reflects the optimal trade-off between high similarity to reference partition and high inter-individual variability in data-driven partitions. *MI*_d−d_ - average mutual information between all pairs of data-driven partitions; *MI_d−r_* - average mutual information between data-driven partitions and reference partition.

#### 2.3.4 Consensus partition composition for various spatial scales

**Table S4.**
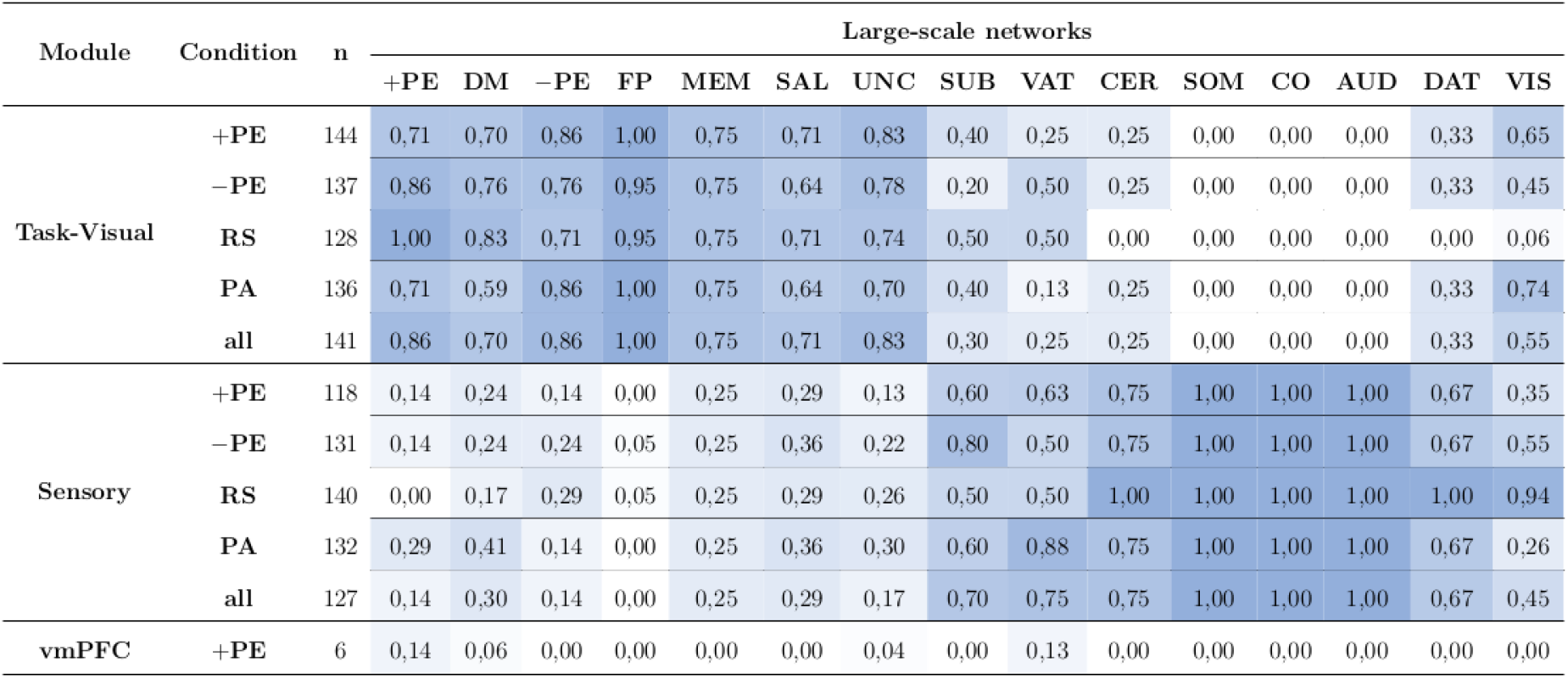
Consensus partition composition for structural resolution parameter γ = 0.5. Cell values correspond to a percentage of reference network regions belonging to the consensus module. Cell color intensity reflects the underlying percentage value. The reference partition consists of 13 well-known LSNs from the Power atlas (Power et al., 2011) and two prediction-error-signaling networks (see section 4.6.1). The third column shows the consensus module size. Task condition abbreviations: +PE - positive prediction error trials, −PE - negative prediction error trials, RS - risk-seeking condition, PA - punishment-avoiding condition. Reference LSNs abbreviations: DM - default mode, FP - fronto-parietal, MEM - memory, SAL - salience, UNC - uncertain, SUB - subcortical, VAT - ventral attention, CER - cerebellar, SOM - somatomotor, CO - cingulo-opercular, AUD - auditory, DAT - dorsal attention, VIS - visual.

**Table S5.** Consensus partition composition for structural resolution parameter γ = 1. Consensus partition composition for structural resolution parameter γ = 1. For abbreviations and additional description see Table S4.

| Module | Condition | n | Large-scale networks |  |  |  |  |  |  |  |  |  |  |  |  |  |  |
| --- | --- | --- | --- | --- | --- | --- | --- | --- | --- | --- | --- | --- | --- | --- | --- | --- | --- |
|  |  |  | +PE | DM | −PE | FP | MEM | SAL | UNC | SUB | VAT | CER | SOM | CO | AUD | DAT | VIS |
| Task-Visual | +PE | 96 | 0,57 | 0,06 | 0,71 | 0,86 | 0,75 | 0,64 | 0,52 | 0,10 | 0,00 | 0,00 | 0,03 | 0,00 | 0,00 | 0,44 | 0,84 |
|  | −PE | 98 | 0,57 | 0,06 | 0,76 | 0,90 | 0,75 | 0,71 | 0,57 | 0,10 | 0,00 | 0,25 | 0,00 | 0,00 | 0,00 | 0,33 | 0,81 |
|  | RS | 87 | 0,29 | 0,02 | 0,67 | 0,86 | 0,75 | 0,57 | 0,48 | 0,10 | 0,00 | 0,00 | 0,03 | 0,00 | 0,00 | 0,33 | 0,81 |
|  | PA | 107 | 0,57 | 0,07 | 0,81 | 1,00 | 0,75 | 0,71 | 0,57 | 0,10 | 0,00 | 0,25 | 0,03 | 0,00 | 0,00 | 0,33 | 0,94 |
|  | all | 94 | 0,57 | 0,04 | 0,71 | 0,86 | 0,75 | 0,71 | 0,48 | 0,10 | 0,00 | 0,00 | 0,03 | 0,00 | 0,00 | 0,33 | 0,84 |
| Default Mode | +PE | 75 | 0,43 | 0,85 | 0,19 | 0,14 | 0,00 | 0,14 | 0,43 | 0,30 | 0,50 | 0,00 | 0,00 | 0,00 | 0,00 | 0,00 | 0,00 |
|  | −PE | 70 | 0,43 | 0,85 | 0,10 | 0,10 | 0,00 | 0,00 | 0,43 | 0,30 | 0,50 | 0,00 | 0,00 | 0,00 | 0,00 | 0,00 | 0,00 |
|  | RS | 76 | 0,71 | 0,87 | 0,19 | 0,14 | 0,00 | 0,14 | 0,43 | 0,10 | 0,50 | 0,00 | 0,00 | 0,00 | 0,00 | 0,00 | 0,00 |
|  | PA | 72 | 0,43 | 0,91 | 0,14 | 0,00 | 0,00 | 0,07 | 0,39 | 0,30 | 0,50 | 0,00 | 0,00 | 0,00 | 0,00 | 0,00 | 0,00 |
|  | all | 74 | 0,43 | 0,87 | 0,14 | 0,14 | 0,00 | 0,07 | 0,43 | 0,30 | 0,50 | 0,00 | 0,00 | 0,00 | 0,00 | 0,00 | 0,00 |
| Sensory | +PE | 97 | 0,00 | 0,09 | 0,10 | 0,00 | 0,25 | 0,21 | 0,04 | 0,60 | 0,50 | 1,00 | 0,97 | 1,00 | 1,00 | 0,56 | 0,16 |
|  | −PE | 100 | 0,00 | 0,09 | 0,14 | 0,00 | 0,25 | 0,29 | 0,00 | 0,60 | 0,50 | 0,75 | 1,00 | 1,00 | 1,00 | 0,67 | 0,19 |
|  | RS | 105 | 0,00 | 0,11 | 0,14 | 0,00 | 0,25 | 0,29 | 0,09 | 0,80 | 0,50 | 1,00 | 0,97 | 1,00 | 1,00 | 0,67 | 0,19 |
|  | PA | 89 | 0,00 | 0,02 | 0,05 | 0,00 | 0,25 | 0,21 | 0,04 | 0,60 | 0,50 | 0,75 | 0,97 | 1,00 | 1,00 | 0,67 | 0,06 |
|  | all | 100 | 0,00 | 0,09 | 0,14 | 0,00 | 0,25 | 0,21 | 0,09 | 0,60 | 0,50 | 1,00 | 0,97 | 1,00 | 1,00 | 0,67 | 0,16 |

## References

Bassett, D. S., and Sporns, O. (2017). Network neuroscience. Nature Neuroscience, 20(3), 353–364.

Bassett, D. S., Porter, M. A., Wymbs, N. F., Grafton, S. T., Carlson, J. M., and Mucha, P. J. (2013). Robust detection of dynamic community structure in networks. Chaos: An Interdisciplinary Journal of Nonlinear Science, 23(1):013142.

Bassett, D. S., Yang, M., Wymbs, N. F., and Grafton, S. T. (2015). Learning-induced autonomy of sensorimotor systems. Nature Neuroscience, 18(5):744–751.

Bavard, S., Lebreton, M., Khamassi, M., Coricelli, G., and Palminteri, S. (2018). Reference-point centering and range-adaptation enhance human reinforcement learning at the cost of irrational preferences. Nature Communications, 9(1):1–12.

Benjamini, Y. and Hochberg, Y. (1995). Controlling the false discovery rate: a practical and powerful approach to multiple testing. Journal of the Royal Statistical Society: Series B (Methodological), 57(1):289–300.

Bertolero, M. A., Yeo, B. T., and D’Esposito, M. (2015). The modular and integrative functional architecture of the human brain. Proceedings of the National Academy of Sciences, 112(49):E6798–E6807.

Betzel, R. F. and Bassett, D. S. (2017). Multi-scale brain networks. Neuroimage, 160:73–83.

Betzel, R. F., Medaglia, J. D., Papadopoulos, L., Baum, G. L., Gur, R., Gur, R., Roalf, D., Satterthwaite, T. D., and Bassett, D. S. (2017). The modular organization of human anatomical brain networks: Accounting for the cost of wiring. Network Neuroscience, 1(1):42–68.

Blondel, V. D., Guillaume, J.-L., Lambiotte, R., and Lefebvre, E. (2008). Fast unfolding of communities in large networks. Journal of Statistical Mechanics: Theory and Experiment, 2008(10):P10008.

Bullmore, E. and Sporns, O. (2009). Complex brain networks: graph theoretical analysis of structural and functional systems. Nature Reviews Neuroscience, 10(3):186–198.

Camara, E., Rodriguez-Fornells, A., and Münte, T. F. (2009). Functional connectivity of reward processing in the brain. Frontiers in Human Neuroscience, 2:19.

Cisek, P. and Kalaska, J. F. (2005). Neural correlates of reaching decisions in dorsal premotor cortex: specification of multiple direction choices and final selection of action. Neuron, 45(5):801–814.

Cisler, J. M., Bush, K., and Steele, J. S. (2014). A comparison of statistical methods for detecting context-modulated functional connectivity in fmri. Neuroimage, 84:1042– 1052.

Cohen, M., Heller, A., and Ranganath, C. (2005). Functional connectivity with anterior cingulate and orbitofrontal cortices during decision-making. Cognitive Brain Research, 23(1):61–70.

Colombo, M. (2014). Deep and beautiful. the reward prediction error hypothesis of dopamine. Studies in history and philosophy of science part C: Studies in history and philosophy of biological and biomedical sciences, 45:57–67.

Conrad, B. N., Wilkey, E. D., Yeo, D. J., and Price, G. R. (2020). Network topology of symbolic and nonsymbolic number comparison. Network Neuroscience, 4(3):714–745.

Cools, R., Clark, L., Owen, A. M., and Robbins, T. W. (2002). Defining the neural mechanisms of probabilistic reversal learning using event-related functional magnetic resonance imaging. Journal of Neuroscience, 22(11):4563–4567.

Daw, N. D., Gershman, S. J., Seymour, B., Dayan, P., and Dolan, R. J. (2011). Model-based influences on humans’ choices and striatal prediction errors. Neuron, 69(6):1204–1215.

Deco, G., Tononi, G., Boly, M., & Kringelbach, M. L. (2015). Rethinking segregation and integration: contributions of whole-brain modelling. Nature Reviews Neuroscience, 16(7), 430–439.

Dehaene, S., Kerszberg, M., and Changeux, J.-P. (1998). A neuronal model of a global workspace in effortful cognitive tasks. Proceedings of the National Academy of Sciences, 95(24):14529–14534.

Doron, K. W., Bassett, D. S., and Gazzaniga, M. S. (2012). Dynamic network structure of interhemispheric coordination. Proceedings of the National Academy of Sciences, 109(46):18661–18668.

Esteban, O., Markiewicz, C. J., Blair, R. W., Moodie, C. A., Isik, A. I., Erramuzpe, A., Kent, J. D., Goncalves, M., DuPre, E., Snyder, M., et al. (2019). fMRIPrep: a robust preprocessing pipeline for functional mri. Nature Methods, 16(1):111–116.

Fazeli, S. and Büchel, C. (2018). Pain-related expectation and prediction error signals in the anterior insula are not related to aversiveness. Journal of Neuroscience, 38(29):6461–6474.

Finc, K., Bonna, K., Lewandowska, M., Wolak, T., Nikadon, J., Dreszer, J., Duch, W., and Kühn, S. (2017). Transition of the functional brain network related to increasing cognitive demands. Human Brain Mapping, 38(7):3659–3674.

Fouragnan, E., Retzler, C., and Philiastides, M. G. (2018). Separate neural representations of prediction error valence and surprise: Evidence from an fMRI meta-analysis. Human Brain Mapping, 39(7):2887–2906.

Frank, M. J. and Claus, E. D. (2006). Anatomy of a decision: striato-orbitofrontal interactions in reinforcement learning, decision making, and reversal. Psychological Review, 113(2):300.

Frank, M. J., Moustafa, A. A., Haughey, H. M., Curran, T., and Hutchison, K. E. (2007). Genetic triple dissociation reveals multiple roles for dopamine in reinforcement learning. Proceedings of the National Academy of Sciences, 104(41):16311–16316.

Freeman, L. C. (1977). A set of measures of centrality based on betweenness. Sociometry, pages 35–41.

Gerraty, R. T., Davidow, J. Y., Foerde, K., Galvan, A., Bassett, D. S., and Shohamy, D. (2018). Dynamic flexibility in striatal-cortical circuits supports reinforcement learning. Journal of Neuroscience, 38(10):2442–2453.

Gershman, S. J. (2015). Do learning rates adapt to the distribution of rewards? Psychonomic Bulletin & Review, 22(5):1320–1327.

Gershman, S. J. (2016). Empirical priors for reinforcement learning models. Journal of Mathematical Psychology, 71:1–6.

Gorgolewski, K., Burns, C. D., Madison, C., Clark, D., Halchenko, Y. O., Waskom, M. L., and Ghosh, S. S. (2011). Nipype: a flexible, lightweight and extensible neuroimaging data processing framework in python. Frontiers in Neuroinformatics, 5:13.

Gorgolewski, K. J., Auer, T., Calhoun, V. D., Craddock, R. C., Das, S., Duff, E. P., Flandin, G., Ghosh, S. S., Glatard, T., Halchenko, Y. O., et al. (2016). The brain imaging data structure, a format for organizing and describing outputs of neuroimaging experiments. Scientific Data, 3(1):1–9.

Gu, S., Satterthwaite, T. D., Medaglia, J. D., Yang, M., Gur, R. E., Gur, R. C., and Bassett, D. S. (2015). Emergence of system roles in normative neurodevelopment. Proceedings of the National Academy of Sciences, 112(44):13681–13686.

Hare, T. A., Camerer, C. F., and Rangel, A. (2009). Self-control in decision-making involves modulation of the vmpfc valuation system. Science, 324(5927):646–648.

Hauser, T. U., Iannaccone, R., Walitza, S., Brandeis, D., and Brem, S. (2015). Cognitive flexibility in adolescence: neural and behavioral mechanisms of reward prediction error processing in adaptive decision making during development. Neuroimage, 104:347–354.

Hayes, D. J., Duncan, N. W., Xu, J., and Northoff, G. (2014). A comparison of neural responses to appetitive and aversive stimuli in humans and other mammals. Neuroscience & Biobehavioral Reviews, 45:350–368.

Horga, G., Maia, T. V., Marsh, R., Hao, X., Xu, D., Duan, Y., Tau, G. Z., Graniello, B., Wang, Z., Kangarlu, A., et al. (2015). Changes in corticostriatal connectivity during reinforcement learning in humans. Human Brain Mapping, 36(2):793–803.

Huckins, J. F., Adeyemo, B., Miezin, F. M., Power, J. D., Gordon, E. M., Laumann, T. O., Heatherton, T. F., Petersen, S. E., and Kelley, W. M. (2019). Reward-related regions form a preferentially coupled system at rest. Human Brain Mapping, 40(2):361–376.

Khaw, M. W., Glimcher, P. W., and Louie, K. (2017). Normalized value coding explains dynamic adaptation in the human valuation process. Proceedings of the National Academy of Sciences, 114(48):12696–12701.

Lancichinetti, A. and Fortunato, S. (2012). Consensus clustering in complex networks. Scientific Reports, 2(1):1–7.

Levy, D. J. and Glimcher, P. W. (2012). The root of all value: a neural common currency for choice. Current Opinion in Neurobiology, 22(6):1027–1038.

Louie, K., Khaw, M. W., and Glimcher, P. W. (2013). Normalization is a general neural mechanism for context-dependent decision making. Proceedings of the National Academy of Sciences, 110(15):6139–6144.

Luce, R. D. (1957). A theory of individual choice behavior. Technical report, Columbia University New York Bureau of Applied Social Research.

Mattar, M. G., Thompson-Schill, S. L., and Bassett, D. S. (2018). The network architecture of value learning. Network Neuroscience, 2(02):128–149.

Meder, D., Madsen, K. H., Hulme, O., and Siebner, H. R. (2016). Chasing probabilities—signaling negative and positive prediction errors across domains. Neuroimage, 134:180–191.

Meder, D., Rabe, F., Morville, T., Madsen, K. H., Koudahl, M. T., Dolan, R. J., Siebner, H. R., and Hulme, O. J. (2019). Ergodicity-breaking reveals time optimal decision making in humans. arXiv preprint arXiv:1906.04652.

Meilă, M. (2007). Comparing clusterings—an information based distance. i, 98(5):873–895.

Montague, P. R., Dayan, P., & Sejnowski, T. J. (1996). A framework for mesencephalic dopamine systems based on predictive Hebbian learning. Journal of Neuroscience, 16(5), 1936–1947.

Mumford, J. A., Turner, B. O., Ashby, F. G., and Poldrack, R. A. (2012). Deconvolving bold activation in event-related designs for multivoxel pattern classification analyses. Neuroimage, 59(3):2636–2643.

Münte, T. F., Heldmann, M., Hinrichs, H., Marco-Pallares, J., Krämer, U. M., Sturm, V., and Heinze, H.-J. (2008). Nucleus accumbens is involved in human action monitoring: evidence from invasive electrophysiological recordings. Frontiers in Human Neuroscience, 2:11.

Namburi, P., Al-Hasani, R., Calhoon, G. G., Bruchas, M. R., and Tye, K. M. (2016). Architectural representation of valence in the limbic system. Neuropsychopharmacology, 41(7):1697–1715.

Nichols, T. E. and Holmes, A. P. (2002). Nonparametric permutation tests for functional neuroimaging: a primer with examples. Human Brain Mapping, 15(1):1–25.

Nilsson, H., Rieskamp, J., and Wagenmakers, E.-J. (2011). Hierarchical bayesian parameter estimation for cumulative prospect theory. Journal of Mathematical Psychology, 55(1):84–93.

Niv, Y., Edlund, J. A., Dayan, P., and O’Doherty, J. P. (2012). Neural prediction errors reveal a risk-sensitive reinforcement-learning process in the human brain. Journal of Neuroscience, 32(2):551–562.

O’Doherty, J. P., Hampton, A., and Kim, H. (2007). Model-based fmri and its application to reward learning and decision making. Annals of the New York Academy of Sciences, 1104(1):35–53.

Palminteri, S., Khamassi, M., Joffily, M., and Coricelli, G. (2015). Contextual modulation of value signals in reward and punishment learning. Nature Communications, 6(1):1–14.

Palminteri, S. and Pessiglione, M. (2017). Opponent brain systems for reward and punishment learning: causal evidence from drug and lesion studies in humans. In Decision neuroscience, pages 291–303. Elsevier.

Peirce, J. W. (2007). Psychopy—psychophysics software in python. Journal of Neuroscience Methods, 162(1-2):8–13.

Power, J. D., Cohen, A. L., Nelson, S. M., Wig, G. S., Barnes, K. A., Church, J. A., Vogel, A. C., Laumann, T. O., Miezin, F. M., Schlaggar, B. L., et al. (2011). Functional network organization of the human brain. Neuron, 72(4):665–678.

Rangel, A., Camerer, C., and Montague, P. R. (2008). A framework for studying the neurobiology of value-based decision making. Nature Reviews Neuroscience, 9(7):545–556.

Rangel, A. & Clithero, J. A. (2012). Value normalization in decision making: theory and evidence. Current Opinion in Neurobiology, 22(6):970–981.

Rao, R. P., and Ballard, D. H. (1999). Predictive coding in the visual cortex: a functional interpretation of some extra-classical receptive-field effects. Nature Neuroscience, 2(1), 79–87.

Reiter, A. M., Koch, S. P., Schröger, E., Hinrichs, H., Heinze, H.-J., Deserno, L., and Schlagenhauf, F. (2016). The feedback-related negativity codes components of abstract inference during reward-based decision-making. Journal of Cognitive Neuroscience, 28(8):1127–1138.

Rescorla, R. A. and Wagner, A. R. (1972). A theory of Pavlovian conditioning: Variations on the effectiveness of reinforcement and non-reinforcement. In Black, A. H. and Prokasy, W. F., editors, Classical conditioning II: Current research and theory, pages 64–99. Appleton-Century-Crofts, New York.

Rigoli, F. (2019). Reference effects on decision-making elicited by previous rewards. Cognition, 192:104034.

Rigoux, L., Stephan, K. E., Friston, K. J., & Daunizeau, J. (2014). Bayesian model selection for group studies—revisited. Neuroimage, 84, 971–985.

Rissman, J., Gazzaley, A., and D’Esposito, M. (2004). Measuring functional connectivity during distinct stages of a cognitive task. Neuroimage, 23(2):752–763.

Robinson, O. J., Frank, M. J., Sahakian, B. J., and Cools, R. (2010). Dissociable responses to punishment in distinct striatal regions during reversal learning. Neuroimage, 51(4), 1459–1467.

Roy, M., Shohamy, D., and Wager, T. D. (2012). Ventromedial prefrontal-subcortical systems and the generation of affective meaning. Trends in Cognitive Sciences, 16(3):147–156.

Rubinov, M. and Sporns, O. (2010). Complex network measures of brain connectivity: uses and interpretations. Neuroimage, 52(3):1059–1069.

Rubinov, M. and Sporns, O. (2011). Weight-conserving characterization of complex functional brain networks. Neuroimage, 56(4):2068–2079.

Sadler, J. R., Shearrer, G. E., Acosta, N. T., Papantoni, A., Cohen, J. R., Small, D. M., Park, S. Q., Gordon-Larsen, P., and Burger, K. S. (2020). Network organization during probabilistic learning via taste outcomes. Physiology & Behavior, 223:112962.

Shine, J. M., Bissett, P. G., Bell, P. T., Koyejo, O., Balsters, J. H., Gorgolewski, K. J., Moodie, C. A., and Poldrack, R. A. (2016). The dynamics of functional brain networks: integrated network states during cognitive task performance. Neuron, 92(2):544–554.

Shteingart, H. and Loewenstein, Y. (2014). Reinforcement learning and human behavior. Current Opinion in Neurobiology, 25:93–98.

Sutton, R. S. and Barto, A. G. (2018). Reinforcement learning: An introduction. MIT press.

Van den Bos, W., Cohen, M. X., Kahnt, T., and Crone, E. A. (2012). Striatum–medial prefrontal cortex connectivity predicts developmental changes in reinforcement learning. Cerebral Cortex, 22(6):1247–1255.

Vatansever, D., Menon, D. K., Manktelow, A. E., Sahakian, B. J., and Stamatakis, E. A. (2015). Default mode dynamics for global functional integration. Journal of Neuroscience, 35(46):15254–15262.

Yacubian, J., Gläscher, J., Schroeder, K., Sommer, T., Braus, D. F., and Büchel, C. (2006). Dissociable systems for gain-and loss-related value predictions and errors of prediction in the human brain. Journal of Neuroscience, 26(37):9530–9537.

## References

Avants, B. B., Epstein, C. L., Grossman, M., and Gee, J. C. (2008). Symmetric diffeomorphic image registration with cross-correlation: evaluating automated labeling of elderly and neurodegenerative brain. Medical Image Analysis, 12(1):26–41.

Cox, R. W. and Hyde, J. S. (1997). Software tools for analysis and visualization of fmri data. NMR in Biomedicine: An International Journal Devoted to the Development and Application of Magnetic Resonance In Vivo, 10(4-5):171–178.

Dale, A. M., Fischl, B., and Sereno, M. I. (1999). Cortical surface-based analysis: I. segmentation and surface reconstruction. Neuroimage, 9(2):179–194.

Fouragnan, E., Retzler, C., and Philiastides, M. G. (2018). Separate neural representations of prediction error valence and surprise: Evidence from an fMRI meta-analysis. Human Brain Mapping, 39(7):2887–2906.

Fortunato, S. and Barthelemy, M. (2007). Resolution limit in community detection. Proceedings of the National Academy of Sciences, 104(1):36–41.

Friston, K., Buechel, C., Fink, G., Morris, J., Rolls, E., and Dolan, R. J. (1997). Psychophysiological and modulatory interactions in neuroimaging. Neuroimage, 6(3):218–229.

Gamerman, D. and Lopes, H. F. (2006). Markov chain Monte Carlo: stochastic simulation for Bayesian inference. CRC Press.

Greve, D. N. and Fischl, B. (2009). Accurate and robust brain image alignment using boundary-based registration. Neuroimage, 48(1):63–72.

Jenkinson, M. (2003). Fast, automated, n-dimensional phase-unwrapping algorithm. Magnetic Resonance in Medicine: An Official Journal of the International Society for Magnetic Resonance in Medicine, 49(1):193–197.

Kass, R. E., Carlin, B. P., Gelman, A., and Neal, R. M. (1998). Markov chain monte carlo in practice: a roundtable discussion. The American Statistician, 52(2):93–100.

Klein, A., Ghosh, S. S., Bao, F. S., Giard, J., Häme, Y., Stavsky, E., Lee, N., Rossa, B., Reuter, M., Chaibub Neto, E., et al. (2017). Mindboggling morphometry of human brains. PLoS Computational Biology, 13(2):e1005350.

Lancichinetti, A., Fortunato, S., and Kertész, J. (2009). Detecting the overlapping and hierarchical community structure in complex networks. New Journal of Physics, 11(3):033015.

Lee, M. D. and Wagenmakers, E.-J. (2014). Bayesian cognitive modeling: A practical course. Cambridge university press.

Reichardt, J. and Bornholdt, S. (2006). Statistical mechanics of community detection. Physical Review E, 74(1):016110.

Satterthwaite, T. D., Elliott, M. A., Gerraty, R. T., Ruparel, K., Loughead, J., Calkins, M. E., Eickhoff, S. B., Hakonarson, H., Gur, R. C., Gur, R. E., et al. (2013). An improved framework for confound regression and filtering for control of motion artifact in the preprocessing of resting-state functional connectivity data. Neuroimage, 64:240–256.

Sporns, O. (2014). Contributions and challenges for network models in cognitive neuroscience. Nature Neuroscience, 17(5):652–660.

Tustison, N. J., Avants, B. B., Cook, P. A., Zheng, Y., Egan, A., Yushkevich, P. A., and Gee, J. C. (2010). N4itk: improved n3 bias correction. IEEE Transactions on Medical Imaging, 29(6):1310–1320.

van Doorn, J., van den Bergh, D., Böhm, U., Dablander, F., Derks, K., Draws, T., Etz, A., Evans, N. J., Gronau, Q. F., Haaf, J. M., et al. (2021). The jasp guidelines for conducting and reporting a bayesian analysis. Psychonomic Bulletin & Review, 28(3):813–826.

Yoon, U., Fonov, V. S., Perusse, D., Evans, A. C., Group, B. D. C., et al. (2009). The effect of template choice on morphometric analysis of pediatric brain data. Neuroimage, 45(3):769–777.

Zhang, Y., Brady, J. M., and Smith, S. (2000). Hidden markov random field model for segmentation of brain MR image. In Medical Imaging 2000: Image Processing, volume 3979, pages 1126–1137. International Society for Optics and Photonics.

